# Competing routes back to firing shape population hyperexcitability and its selective suppression

**DOI:** 10.64898/2026.11.09.765141

**Authors:** Aravind Kumar Kamaraj, Matthew Parker Szuromi

## Abstract

Even within the same neuron, successive spikes can follow distinct trajectories, with afterdepolarization permitting early return to firing and depolarization block prolonging inactivity. Therefore, we propose that hyperexcitability involves changes in the opportunities to fire again from different points across the spike cycle, and not just from quiescence. The *hazard of firing* — the instantaneous propensity to fire again with time elapsed after a spike — isolates these opportunities. In bursting dopamine neurons, the hazard rises early due to burst-sustaining spikes, falls, then rises again due to burst-initiating spikes. Within a renewal framework, we represent each rise as a competing pathway back to firing and show that recruitment from quiescence and firing again before full recovery provide independent control over population activity. Changes in pathway engagement distinguish subclinical from overt hyperexcitability and allow neurons of the same type to respond differently to a common input. Adding a slower return pathway for drug-bound neurons shows how sodium-channel blockers such as phenytoin and carbamazepine can suppress pathological bursting while leaving physiological activity apparently intact.

## Introduction

Neuronal hyperexcitability has been implicated as a pathophysiological hallmark across a remarkably diverse set of neurological disorders, including epilepsy [1, 2], Alzheimer’s disease [3, 4], amyotrophic lateral sclerosis (ALS) [5, 6], and chronic pain [7, 8]. Yet the field has never settled on a uniform, mechanism-based definition. At the population level, hyperex-citability can arise from more neurons firing, or from the same neurons firing more often. The first is a question of recruitment — how many neurons are drawn into activity. The second is a question of recurrence — how readily a neuron that has just fired can fire again, which is set by the route it takes back to firing. In existing accounts of hyperexcitability, that route is held fixed and pathology is represented by changes in the rates of the processes along it. But why would a pathology preserve all neuronal processes while altering only their rates? Here, we let pathology bypass some processes altogether, allowing firing to resume by different routes. We then show that hyperexcitability and its selective suppression both follow from a shift in competition among those routes.

Considering neuronal firing as a renewal process, the quantity of interest is the *hazard* of firing: given the time elapsed since the last spike, what is the probability that a neuron that has not fired up to this point would do so now? By retaining the temporal structure of firing propensity, the hazard characterizes excitability more faithfully than a scalar metric. For instance, in bursting neurons, clustered firing produces a bimodal hazard, where spikes within a burst give rise to an early peak within about 10 ms of a spike; the hazard then declines over the following tens of milliseconds, and burst initiating spikes lead to a later rise [9, 10]. Indeed, hazards of both monotone and non-monotone form have long been examined at the level of single neurons [11, 12].

Hyperexcitability, therefore, need not increase the firing propensity uniformly across the spike cycle but may instead selectively amplify particular opportunities to fire again. NMDA receptor conductance, dysregulated in epilepsy [13] and Alzheimer’s disease [14], illustrates the point. After a spike, the brief depolarizing afterpotential can enhance the NMDA current by relieving its magnesium block, increasing the likelihood of renewed firing early in the spike cycle [15, 16]. Hyperpolarization restores the block and suppresses the NMDA current [17]. Altered NMDA signaling thus provides a shortcut to firing directly from afterdepolarization, introducing an early peak in the firing hazard or amplifying one already present during physiological bursting.

In standard neural mass models of hyperexcitability, such as the Wilson–Cowan framework and its descendants, recovery proceeds at constant rates, and upon reaching quiescence, the activation function governs the rate of return to firing [18, 19]. Pathology, in these models, is then encoded by modifying this activation function [20–23]. Refractory-density [24, 25] and renewal [26, 27] formulations generalize this by making recovery itself input-dependent. These changes usually alter the rates of transition while preserving a single recovery trajectory, and this topology places a restriction: under a fixed input, the firing propensity increases as recovery advances, leading to a monotonically increasing hazard [28]. The population-level consequences of relaxing that topology remain unquantified.

A second line of evidence comes from pharmacology. Anti-seizure medications such as phenytoin and carbamazepine selectively suppress pathological firing while sparing normal physiological function [29, 30]. These drugs bind to inactivated sodium channels in recently active neurons and slow their recovery [31]. The difference in recovery rates between bound and unbound neurons is harder to represent through a single monotone hazard, since it averages over drug-bound and drug-free recovery. Distinct return pathways with well-separated timescales naturally resolve the two.

Here, we represent the various pathways to return to firing as competing hazards, and show how their relative engagement shapes population hyperexcitability and its selective suppression. We first test this premise in *in vivo* whole-cell recordings of midbrain dopamine neurons [32].

In a subset of neurons, the voltage trough preceding each spike falls into two classes, and partitioning interspike intervals by trough class resolves the pooled hazard into components with distinct return dynamics. Representing these components as competing pathways in a continuous-time Markov-chain model yields three results. An *immediate-return pathway*, which bypasses later stages of recovery, controls population activity independently of recruitment and separates signatures of subclinical hyperexcitability from overt seizures. A *sequestration pathway*, representing use-dependent sodium-channel blockade, suppresses pathological activity, because fast-cycling neurons renew, and are therefore sequestered, more often. Finally, competing pathways cause neurons of the same type under a common drive to diverge, accounting for heterogeneous responses at seizure onset that have so far eluded explanation [33, 34].

## Results

### Physiological bursting exhibits a non-monotone hazard with pathway-specific components

Our framework rests on a single structural premise: that a neuron returns to firing by more than one route, rather than by one sequence traversed at varying speed. Before asking what such routing would imply for population activity, we ask whether it can be seen. During bursting, a neuron with the same intrinsic properties generates spikes that are not equivalent: the first spike of a burst follows a deep after-hyperpolarization; subsequent spikes within the burst are preceded by an after-depolarization, and the interspike interval (ISI) differs accordingly — long between bursts, short within them [35, 36].

We examined this structure using the in vivo whole-cell recordings of midbrain dopamine neurons reported by Otomo et al. [32]. The dataset comprises 110 spontaneously active, immunocytochemically identified neurons and is well-suited to this analysis for two reasons. First, the physiological setting reduces the confounding influence of strongly nonstationary population input, supporting a more direct attribution of a non-monotone hazard to competing return pathways [37]. In seizure recordings, by contrast, synchronous population activity generates synaptic input that varies strongly on timescales comparable to ISIs and so it would be difficult to distinguish pathway-specific effects from those of non-stationary input. Second, the subthreshold voltage trajectory from the intracellular recordings provides an observable marker of the state from which firing resumes rather than requiring that state to be inferred from ISIs alone. For each spike, we consider the trough voltage (*V*_min_), defined as the most hyperpolarized potential reached preceding that spike, to represent the intrinsic state from which firing resumed. Of the 110 neurons recorded, 97 had at least 80 resolvable ISIs and were retained for further analysis.

In a neuron exhibiting regular pacemaking (Fig. 1(a)), variation in the ongoing drive produces a single dispersed cluster in log_10_(ISI) (Fig. 1(b)). The intrinsic neuronal processes preceding successive spikes likewise produce a unimodal distribution of *V*_min_. The joint distribution of *V*_min_ and log_10_(ISI) generally exhibits a negative correlation, but the magnitude of this correlation, measured using Spearman’s rank coefficient, varies widely among pacemaking neurons (Fig. 1(k)).

**Fig. 1:**
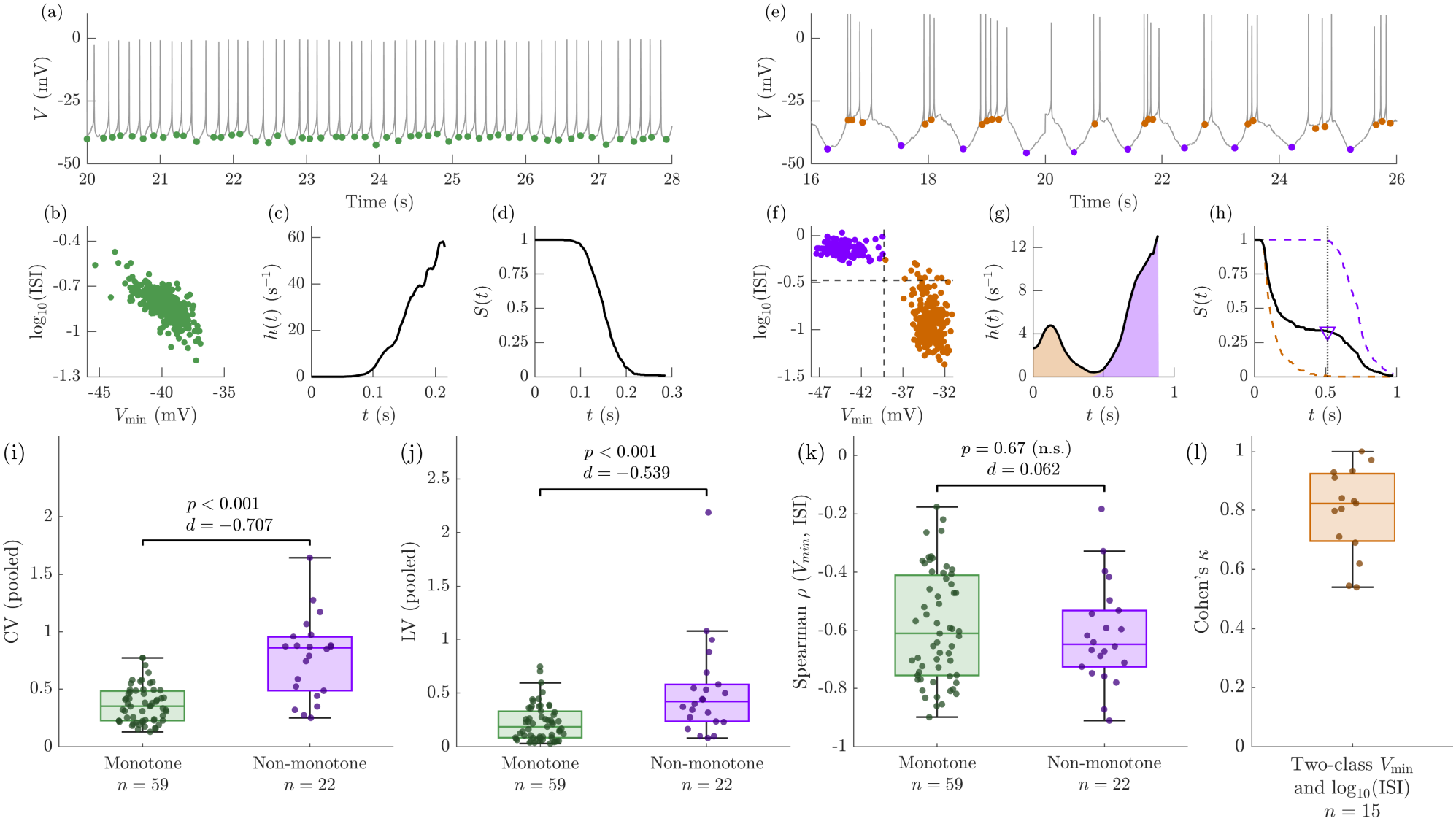
In some neurons, the trough voltage (*V*_min_) reached before a spike admits two classes, each governed by a distinct firing hazard. *(a to d) Representative neuron exhibiting regular pacemaker firing*. (a) Time course of membrane potential (*V* ), with *V*_min_ preceding each spike marked in green. (b) The joint distribution of *V*_min_ and log_10_(ISI) forms a single cluster, consistent with repeated return to firing from one post-spike state. (c) The corresponding hazard remains monotone increasing. (d) The associated survival function. *(e to h) Representative neuron exhibiting burst firing*. (e) Burst-initiating spikes follow deep hyperpolarized troughs (purple), whereas subsequent intraburst spikes follow more depolarized troughs (orange). (f) The joint distribution of *V*_min_ and log_10_(ISI) resolves two clusters, showing that each *V*_min_ class is associated with a distinct ISI class. Each point represents one spike, colored based on the two *V*_min_ classes. (g) The corresponding hazard is non-monotone, admitting a two-component mixture. (h) The corresponding survival function and its two class-specific components. *(i to l) Group comparisons*. (i) Coefficient of variation and (j) local variation are significantly higher in neurons with non-monotone hazard compared to those with monotone hazard. (k) Spearman rank correlation between *V*_min_ and log_10_(ISI) per neuron does not show significant differences between the two groups. (l) Classifications obtained independently from *V*_min_ and log_10_(ISI), quantified using Cohen’s *κ*, indicate near-perfect agreement (median ≈ 0.82) in neurons that admit a two-class representation of both quantities. Recordings were sourced from Otomo et al. [32]. Significance thresholds are based on the Mann-Whitney U-test [38], and the associated effect sizes are estimated non-parametrically using Cliff’s delta [39].

A different organization was evident in bursting neurons (Fig. 1(e)), where either *V*_min_ or log_10_(ISI) or both, separated into two classes: burst-initiating spikes were preceded by deep hyperpolarization, whereas subsequent intraburst spikes arose from relatively depolarized troughs. For each neuron, we independently applied kernel density estimation or if that failed, a shoulder criterion to identify the classes of the *V*_min_ and the log_10_(ISI) distributions (see Methods for the classification procedure).

Plotting *V*_min_ against the corresponding ISI showed that the hyperpolarized class was associated predominantly with longer intervals, and the depolarized class with shorter intervals. In Fig. 1(f), dashed vertical and horizontal lines indicate the class boundaries in *V*_min_ and log_10_(ISI), respectively. The limited overlap between these independently resolved classifications suggests that *V*_min_ contains information about the timing of the following spike.

To formalize this relationship, let *T*_*F F*_ > 0 denote the ISI, with probability density *f* (*t*). As the occurrence of the next spike terminates the interval, the survival function — the probability that the neuron has not fired by time *t* after its preceding spike — is

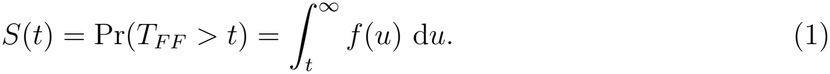

The corresponding hazard function gives the instantaneous firing rate at time *t*, conditional on survival without firing until that time:

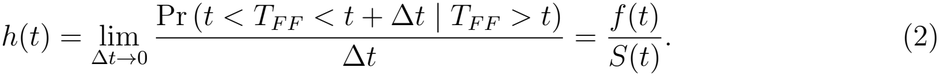

For each neuron, we estimated *S*(*t*) = Pr(ISI > *t*) using the empirical survival function and the hazard by kernel smoothing. The two-class log_10_(ISI) distributions observed in bursting neurons are associated with a non-monotonic dependence of firing propensity on the time elapsed since the preceding spike: the hazard initially rises, subsequently falls, and then rises again. Therefore, we formally tested for departures from hazard monotonicity using the isotonic projection test [40], classifying 63 neurons as monotone and 34 as non-monotone; representative neurons for each category are shown in Figs. 1(c,d) and Figs. 1(g,h), respectively.

Among the 34 neurons with non-monotone hazards, 23 admitted two classes in *V*_min_, 20 in log_10_(ISI), and 15 in both. In each, the first *V*_min_ class was assigned to the more hyperpolarized state. We then partitioned the intervals by the identified *V*_min_ classes and estimated the class-specific survival and hazard functions using the same procedures applied to the pooled intervals. The aggregate hazard then decomposes as,

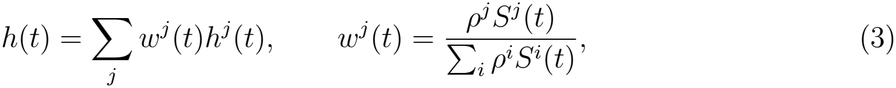

where *ρ*^*j*^ is the mixing proportion of class *j*, and *S*^*j*^(*t*) and *h*^*j*^(*t*) are its class-specific survival and hazard functions, respectively. The weights *w*^*j*^(*t*) are the posterior probabilities that an interval surviving to age *t* belongs to class *j*.

Equation (3) provides a test to assess whether the hazard decomposition represents physiologically distinct classes. If the identified *V*_min_ classes have distinct ISI distributions, their survival functions differ, the weights *w*^*j*^(*t*) vary with time, and the pooled hazard cannot generally be recovered from the fixed-weight combination Σ_*j*_ *ρ*^*j*^*h*^*j*^(*t*). By contrast, under the null hypothesis that the classes are indistinguishable, their survival functions coincide and *w*^*j*^(*t*) = *ρ*^*j*^ for all *t*. We evaluated this null by permuting the class labels while preserving class sizes. The null was rejected in 22 of the 23 neurons with two fitted *V*_min_ classes (200 permutations; *p* < 0.025), demonstrating that the corresponding classes had distinct interval distributions. These 22 neurons formed the non-monotone hazard group for subsequent comparisons; remaining neurons initially classified as non-monotone were excluded from group-level analyses. Of the 63 neurons initially classified as having monotone hazards, four showed bimodal *V*_min_ distributions and were likewise excluded, leaving 59 neurons in the monotone hazard group. Analyses of excluded neurons are reported in the Supplementary Information.

For these two groups, we quantified firing variability using the coefficient of variation (CV) and the local variation (LV) measure introduced by Shinomoto et al. [41]. Unlike CV, LV compares successive ISIs and is therefore less sensitive to slow non-stationarity in the firing rate. Both CV and LV were significantly greater in the non-monotone hazard group than in the monotone hazard group, with large effect sizes (Figs. 1(i) and 1(j)), consistent with the coexistence of short intraburst and long interburst timescales.

Several observations support the interpretation that the two *V*_min_ classes represent distinct post-spike recovery pathways. Among the 15 neurons with independently fitted two-class representations of both *V*_min_ and log_10_(ISI), the median Cohen’s *κ* was approximately 0.82, indicating almost perfect agreement between voltage-defined and ISI-defined classes. By contrast, the within-cell Spearman correlation between *V*_min_ and log_10_(ISI) varied broadly, from approximately −0.2 to −0.9, and did not differ significantly between the monotone and non-monotone hazard groups. The principal distinction between the groups therefore lies not in the continuous co-variation of trough voltage and interval duration, but in the separation of these quantities into discrete, mutually informative classes. Moreover, in 22 of the 23 neurons with a non-monotone hazard and two-class *V*_min_, the voltage-defined classes had interval distributions that were distinguishable from those obtained under label permutation, proving that these classes are governed by distinct firing hazards.

Importantly, these recordings were obtained from spontaneously active dopamine neurons under anesthesia and do not constitute a disease model. Rather, they establish that non-monotone hazards occur under physiological conditions and can be resolved using a directly observed physiological variable. We next use mathematical modeling to examine how pathological hyperexcitability manifests under monotone versus non-monotone hazards.

### A competing hazards framework for neuronal firing

We represent the distinct hazard classes identified above as distinct pathways by which a neuron returns to firing. This pathway structure can be viewed at two levels: within a neuron, different pathways are sampled across successive intervals; across an ensemble of neurons sharing a type and input, different neurons sample different pathways at any given instant.

We model each neuron as a continuous-time Markov chain (CTMC) — a stochastic process that jumps between a discrete set of internal states at continuous times. We consider a homo-geneous population of *N* such neurons, and invoke the mean-field approximation, so that all neurons experience the same effective input, *z*(*t*).

We start with a single, fixed recovery sequence — the canonical pathway — representative of the topological structure underlying conventional models of hyperexcitability: after firing *F*, each neuron traverses a series of intermediate post-spike recovery states *C*_1_, …, *C*_*n*_, left deliberately general, at constant rates *γ*_0_, *γ*_1_, …, *γ*_*n*_. Upon reaching quiescence *Q*, the activation function *A*(*z*) governs the recruitment onto *F* .

There are two ways to represent pathological perturbations to this sequence. The first would be to consider that the pathology modifies the activation *A*(*z*) or the recovery rates *γ*_0_, *γ*_1_, …, *γ*_*n*_ or both. This implicitly assumes that pathology preserves all recovery processes, altering only their rates, with each neuron still having only one path back to *F* .

However, preceding analyses revealed that even physiological neurons can exhibit two distinct spiking windows, captured by a non-monotone hazard. Therefore, a second, broader account of representing pathology is necessary, which involves letting a neuron skip a portion of the canonical recovery sequence and return to firing early from an intermediate state. We implement this through an immediate-return pathway, represented by an edge from *C*_*k*_ to *F* with inputdependent rate *β*_*k*_*ℬ*_*k*_(*z*) (illustrated in Fig. 2 in magenta). Lower values of *k* could represent bursting triggered from after-depolarization, while higher values could represent tonic firing triggered from relative refractoriness.

**Fig. 2:**
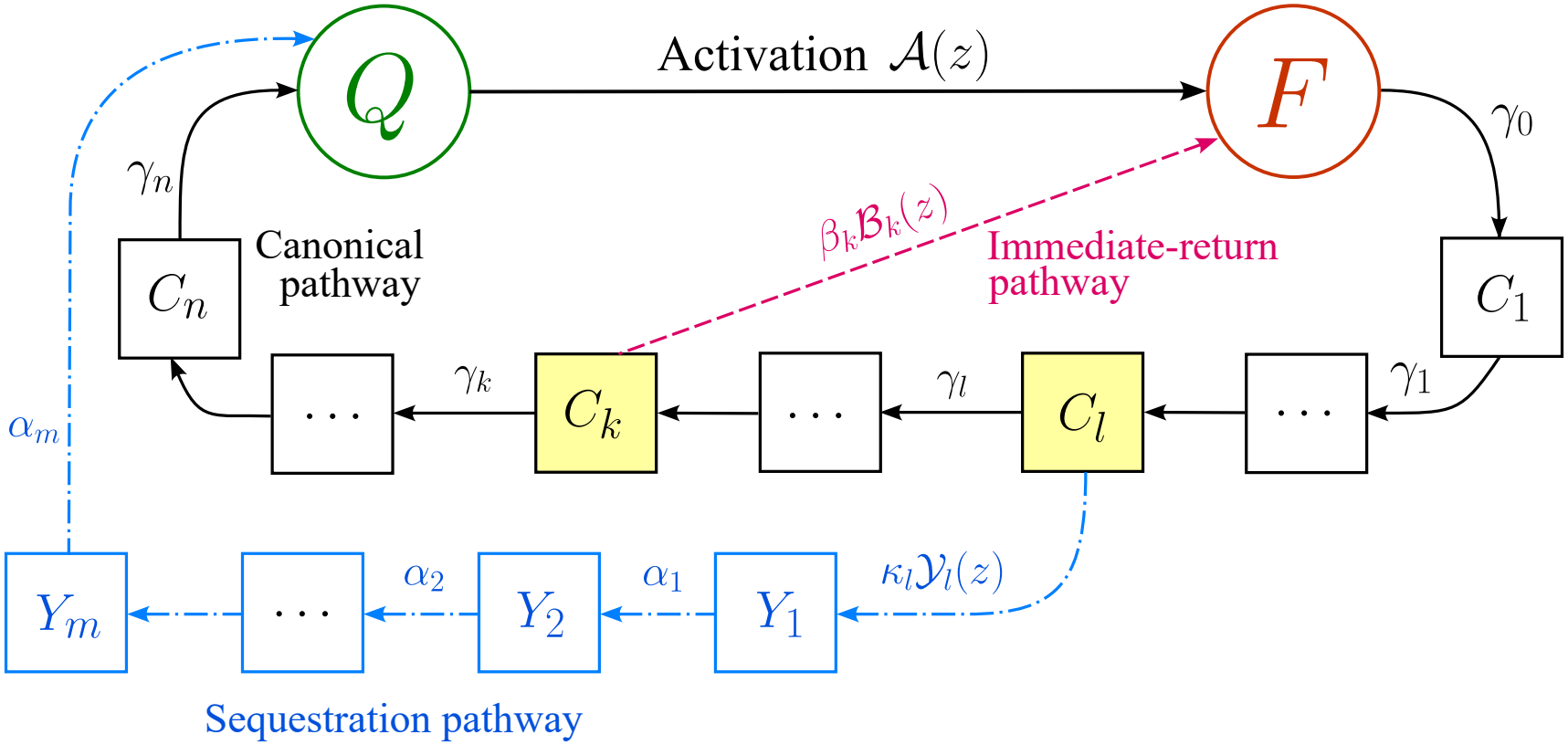
Competing hazards framework for hyperexcitability with canonical, immediatereturn, and sequestration pathways. The canonical pathway is the typical trajectory neurons take: after firing (*F* ), neurons proceed through a series of intermediate post-spike recovery states *C*_1_, …, *C*_*n*_ at constant rates, eventually reaching quiescence (*Q*). Subsequently, the activation function A(*z*) governs the recruitment from *Q* to *F* . We add two kinds of pathways: (i) an immediate-return pathway, the edge denoted in pink, to allow neurons to return to firing from an intermediate state *C*_*k*_ at an input-dependent rate *β*_*k*_B_*k*_(*z*). The input-dependent component B_*k*_(*z*) encodes the observed physiology: different inputs are required to trigger firing from quiescence or a relative refractory state or an after-depolarized state; (ii) a sequestration pathway, the edge denoted in blue, that departs from *C*_*l*_, traverses *m* non-firing, non-recruitable stages with exit rates *α*_1_, *α*_2_, …, *α*_*m*_, and terminates at the quiescent state *Q*. Neurons enter sequestration at an input-dependent rate *κ*_*l*_Y_*l*_(*z*). This pathway could represent depolarization block or the action of anti-seizure medications such as phenytoin and carbamazepine, which preferentially bind to inactivated sodium channels in recently active neurons. The locations *k* and *l* are arbitrary, including their relative placement, and may even coincide (*k* = *l*).

Just as the recovery sequence can be shortened, it can also be prolonged: we add a sequestration pathway that branches off from state *C*_*l*_, traverses *m* non-firing, non-recruitable stages, *Y*_1_, *Y*_2_, … *Y*_*m*_ with exit rates *α*_1_, *α*_2_, …, *α*_*m*_, and terminates at the quiescent state *Q* as shown in Fig. 2. By construction, the mean residence time in the sequestration pathway is strictly greater than the mean residence time of the canonical sequence from *C*_*l*_ to *Q* that it replaces. This pathway could represent depolarization block, where some neurons cease firing at very high levels of input [42], or the action of anti-seizure medications like phenytoin and carbamazepine, which preferentially bind to inactivated sodium channels in recently active neurons and prolong their recovery.

Given an initial condition, the stochastic evolution of each neuron is fully specified by the Markov process generator *G*(*z*), which contains the transition rates between states. The population dynamics can then be expressed in terms of the occupancies — the proportion of neurons in each state — using the Kolmogorov forward equation,

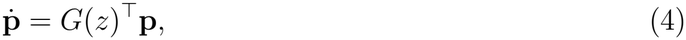

where **p** = (*p*_*s*_)_*s*∈*S*_ is the occupancy of each state *s*, and *z*(*t*) = *ap*_*F*_ + *u*(*t*). These occupancies satisfy the conservation condition,

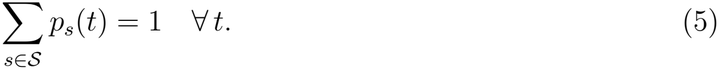

For a full derivation, see Methods.

For characterizing hyperexcitability, the macroscopic quantity of interest is the firing occupancy *p*_*F*_ . The microscopic quantity governing it is the inter-spike interval *T*_*F F*_, which is distributed according to *f* (*z, t*). Here, *T*_*F F*_ is a mixture over three structurally distinct pathways (denoted by superscripts hereafter) — canonical *C*, immediate-return *R*, and sequestration *S*. Therefore, the first return time distribution *f* (*z, t*) decomposes as:

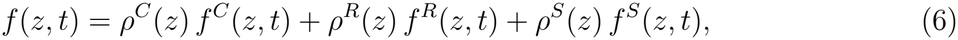

where *ρ*^*j*^(*z*) denotes the probability of returning to firing along pathway *j* and *f*^*j*^(*z, t*) denotes the associated pathway-specific return time distribution. See Methods for the closed-form expressions of *f* (*z, t*) and its mean *µ*(*z*), including pathway-specific contributions. The survival and hazard functions are then evaluated from *f* (*z, t*) using Eqs. (1) and (2) respectively.

We connect *f* (*z, t*) and *p*_*F*_ using a simple renewal framework by restricting attention to steady states, **p**(*t*) = **p**^*^ ∀ *t*. Hence, the self-generated input,

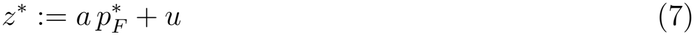

is also constant, rendering the single-neuron CTMC time-homogeneous and the renewal machinery above exact [43].

By the renewal-reward theorem, the steady state firing occupancy is the ratio of the mean occupancy time in *F*, 1*/γ*_0_, to the mean first-return time *µ*(*z*^*^):

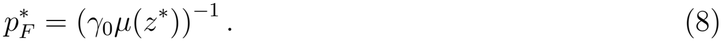

A self-consistent steady state is any pair 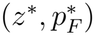 satisfying Eqs. (7) and (8) simultaneously. Since Eq. (8) is nonlinear, multiple self-consistent pairs may exist for the same parameters and external drive *u*, allowing the population to sustain multistability.

### A single pathway back to firing preserves a monotone hazard for all perturbations to parameters

We first restrict the model to consist of only the canonical recovery pathway and examine how varying each parameter associated with this pathway affects the population response. As an illustration, Fig. 3 is generated using a canonical sequence with ten intermediate states (*n* = 10), with equal exit rates *γ*_1_ = *γ*_2_ = · · · = *γ*_*n*_ = *γ*. For each parameter, we plot the 1-D bifurcation diagram between the firing occupancy 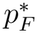 and the external drive *u* in the left column of Fig. 3. The steady state response, being modeled through a 1-D simple renewal framework, only admits fixed points as solutions. We consider the low-activity and high-activity *attractors* (stable fixed points) to be indicative of normal physiological activity and an overt hyperexcitable state such as a seizure, respectively. For a subset of parameter values, the model admits bistability, with co-existing low- and high-activity attractors (solid curves) separated by an unstable fixed point (dotted curve). Two saddle node bifurcations, where a stable and unstable branch collide and annihilate mark the boundaries of the bistable regime. Where bistability is not observed, the model transitions smoothly from low-activity to high-activity as *u* is varied. For select points within this bifurcation diagram, we also plot the first return time distribution and the associated hazard function in the middle and right columns, respectively.

**Fig. 3:**
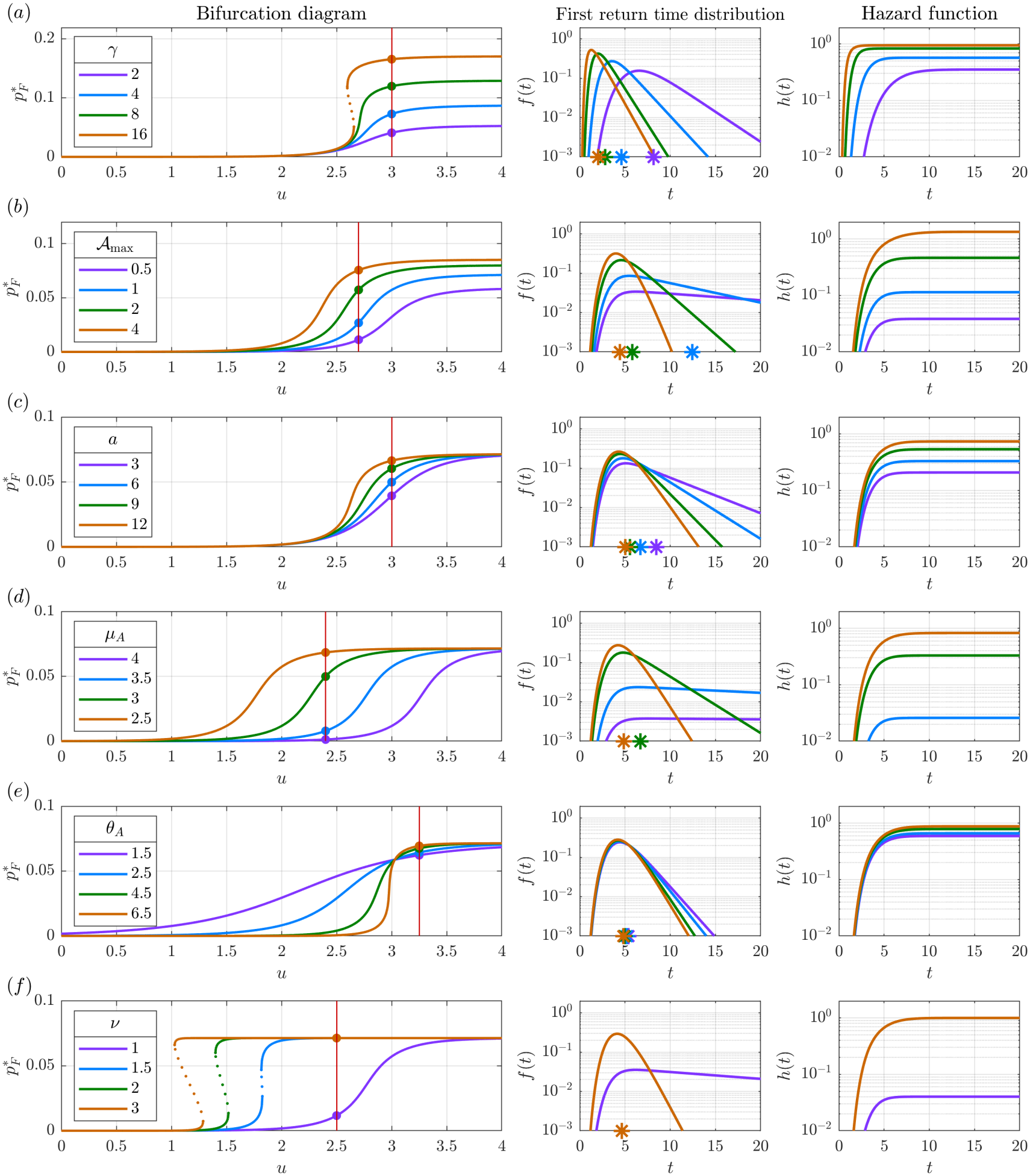
Mechanistically distinct routes to hyperexcitability in the canonical restriction all converge on an upward or leftward shift of a monotonically increasing hazard function. Variation in six different parameters of the canonical restriction lead from a monostable low activity fixed point to a monostable high activity fixed point, with an intervening bistable regime for a subset of parameter values: (a) Increasing recovery rates *γ*_1_ = *γ*_2_ = · · · = *γ*_*n*_ = *γ*, (b) Increasing the supremum of the activation function A_max_, (c) Increasing the mean synaptic weight *a*, (d) Decreasing the activation threshold *µ*_*A*_, (e) Increasing the activation slope *θ*_*A*_ and (f) Increasing the degree of superlinearity of the input *ν*. In each case, the left panels show the 1-D bifurcation diagram for the variation in the firing occupancy 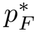 against external drive *u*. Stable branches are indicated in solid lines while the unstable branch is indicated in dotted line. The middle and the right panels show the first return time distribution *f* (*t*) (with its mean indicated as an asterisk) and the hazard function *h*(*t*) at select points indicated in the bifurcation diagram. In all cases, the hazard remains monotone increasing in response to parameter perturbations while the firing occupancy saturates at high levels of external drive. Modifying the recovery rates or the supremum A_max_ alters this saturation ceiling (Panels a and b), while holding them constant and varying the remaining activation parameters leads to the occupancy saturating at the same ceiling (Panels c to f). Parameters are set as in Supplementary Material.

Increasing *γ* selectively increases the level of activity sustained in the high-activity attractor while having minimal influence on the low-activity attractor. Very high values of *γ* lead to bistability. For each value of *γ*, the level of sustained activity saturates with an increase in *u*, with this saturation ceiling itself increasing with an increase in *γ*. Holding *γ* constant, and increasing *A*_max_ leads to a similar response, but the influence is exerted on both low- and high-activity attractors.

In the return time distributions plotted in Fig. 3(b), for the same level of external drive *u*, a higher activation leads to neurons cycling an order of magnitude faster: *f* (*t*) shows an earlier, sharper peak, and faster decline with time *t* during high-activity compared to low-activity. Therefore, during low activation, the term 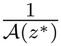 acts as the bottleneck governing the rate of return and so increasing *A*_max_ increases the activity levels, while changing *γ* has little effect. On the other hand, during high activation, *A*(*z*^*^) approaches *A*_max_, and the latter along with *γ* becomes the bottleneck, and increasing either therefore lifts the saturation ceiling.

Varying other parameters of the activation function *A*(*z*), such as its threshold *µ*_*A*_ and slope *θ*_*A*_, and those of its argument, such as the mean synaptic weight *a*, govern the range of *u* over which each attractor exists, but these parameters do not influence the saturation ceiling by themselves. They all converge to the same ceiling as *u* increases (Fig. 3(c) to (e)), as the activation function saturates to its maximum permissible value for high values of 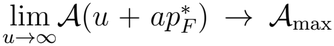. A similar behavior is observed when the input to the activation function is scaled by a factor *ν, A*((*z*^*^)^*ν*^), making it superlinear [22] (Fig. 3(f)).

Importantly, at steady state, irrespective of any perturbation to any parameter, the hazard of firing always remains monotone increasing with time (right column of Fig. 3), and asymptotically approaches the slowest transition rate [43],

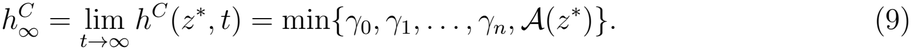

For proof, see Theorem 1 in Methods. In each case, an increase in 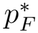 is always accompanied by a leftward or upward shift in the hazard function.

### Differential influence of pathology on immediate-return and canonical pathways dissociates signatures of overt versus subclinical hyperexcitability

We now examine how more than one pathway back to firing influences pathological hyper-excitability by adding an immediate-return pathway from *C*_*k*_ to *F* . Here, the rate of immediate-return *β*_*k*_ quantifies the propensity to take the shorter path back to firing. Increasing *β*_*k*_, with *k* = 2 and all other parameters held fixed, moves the system from a low-activity attractor representative of physiological activity to a high-activity attractor representative of pathological bursting, with bistability observed for an intermediate range of *β*_*k*_ (Fig. 4(a)). In the high-activity regime, increasing *β*_*k*_ progressively increases the firing occupancy, with saturation occurring at very high levels, beyond those explored here.

**Fig. 4:**
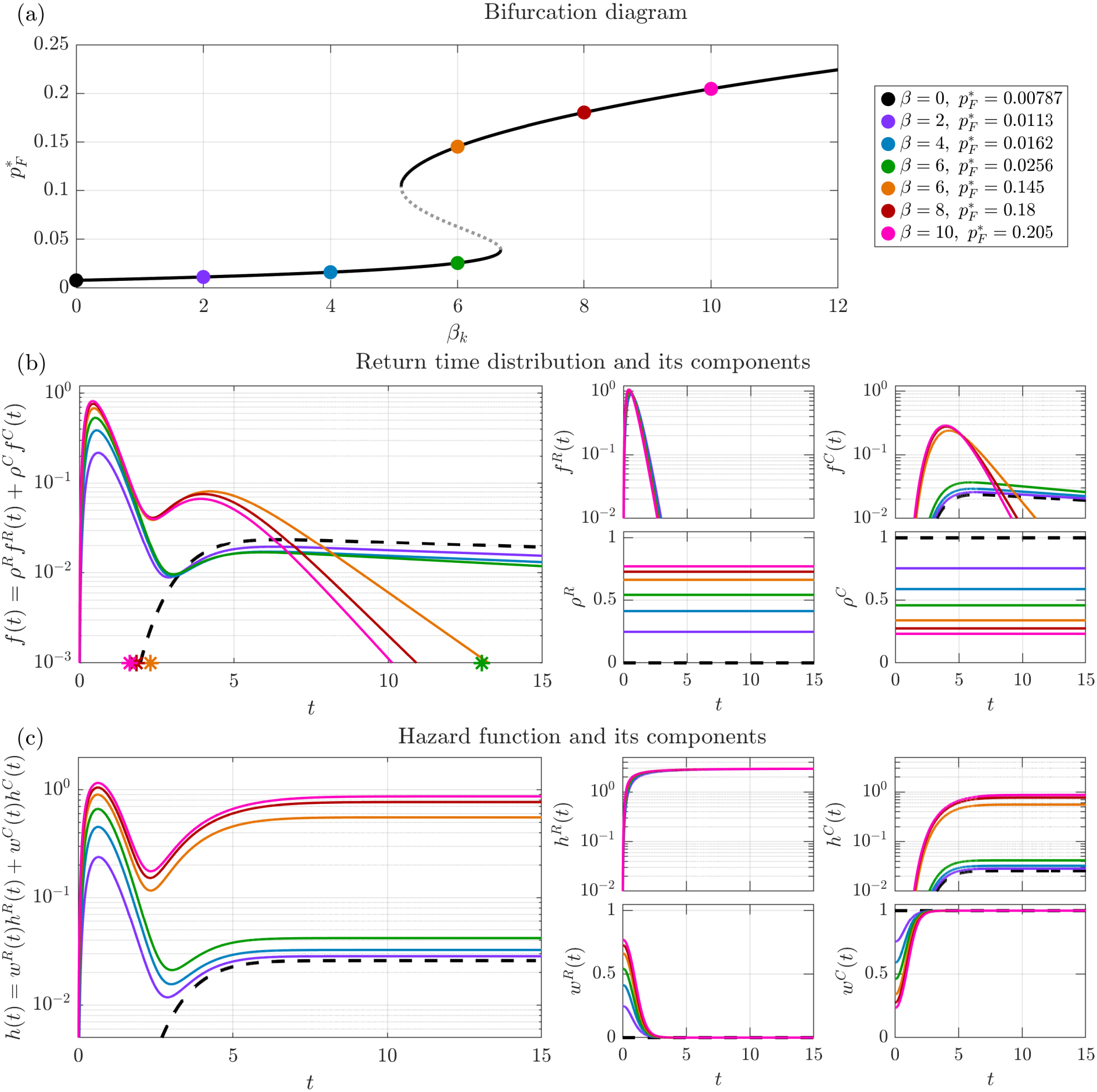
Differential influence of pathological hyperexcitability on immediate-return and canonical pathways. (a) Bifurcation diagram plotting firing occupancy 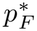 against the rate of immediate-return *β*_*k*_, with *k* = 2 and all other parameters held fixed. Stable branches are indicated in solid lines while the unstable branch is indicated in dotted line. (b) First return time distribution and its components for select points marked in the bifurcation diagram, colored accordingly. The case with the canonical pathway alone (*β*_*k*_ = 0) is shown as black dashed line. For each case, the mean return time is indicated as an asterisk. Note that the mean return times for *β*_*k*_ = 0, 2, and 4 are outside the view of the plot. Here, the component weights *ρ*^*R*^ and *ρ*^*C*^ indicate the probability of returning to firing along immediate-return and canonical pathways, respectively, and remain constant with time elapsed since the last spike. (c) The corresponding hazard function and its components. Here, the component weights *w*^*R*^(*t*) and *w*^*C*^(*t*) indicate the posterior probabilities that a neuron surviving to age *t* returns to firing along immediate-return and canonical pathways, respectively. Immediate-return weights decrease with time since the last spike while canonical weights increase. Therefore, while both hazard components *h*^*R*^(*t*) and *h*^*C*^(*t*) remain monotone increasing, their weighted sum results in a non-monotone aggregate hazard function. Parameters are set as in Supplementary Material.

The canonical and immediate-return pathways are each represented as a series of stages with exponential rates. Therefore, the hazard of firing along each pathway, *h*^*R*^(*t*) and *h*^*C*^(*t*), remains monotone increasing. The corresponding weights, *w*^*R*^(*t*) and *w*^*C*^(*t*), describe the posterior probability of returning along pathway *j* conditional on not having fired by time *t* elapsed since the last spike (Eq. (3)). Immediate-returns occur predominantly at short intervals, progressively depleting this component from the surviving population. Consequently, *w*^*R*^(*t*) monotonically decreases with *t* while *w*^*C*^(*t*) increases, and the aggregate hazard, *h*(*t*) = *w*^*R*^(*t*)*h*^*R*^(*t*)+*w*^*C*^(*t*)*h*^*C*^(*t*), becomes non-monotone (Fig. 4(c)).

On the other hand, the first return time distribution decomposes as *f* (*t*) = *ρ*^*R*^*f* ^*R*^(*t*)+*ρ*^*C*^*f* ^*C*^(*t*), where the mixture weights *ρ*^*R*^ and *ρ*^*C*^ remain constant with *t* (Fig. 4(b)). The weights *ρ*^*R*^ and *ρ*^*C*^ represent the probabilities of returning to firing along immediate-return and canonical pathways, respectively. Both pathways contribute to the low- and high-activity attractors when *β*_*k*_ > 0, and neither physiological activity nor pathology is uniquely attributable to either pathway.

A closer look at the decomposition of *f* (*t*) reveals the role played by each pathway and helps dissociate signatures of overt versus subclinical pathological states. We consider the high-activity attractor to represent a state of overt hyperexcitability like a seizure, and an increase in *β*_*k*_ within the low-activity branch to represent a state of enduring but subclinical hyper-excitability such as those observed during interictal periods or in conditions like Alzheimer’s disease. Furthermore, since immediate-return is placed early in the spike cycle (*k* = 2, *n* = 10), we consider the returns to firing via the canonical pathway as spikes that initiate a burst, and those via immediate-return as successive spikes within a burst.

Within either attractor, increasing *β*_*k*_ increases *ρ*^*R*^ and reduces *ρ*^*C*^, reflecting a higher fraction of returns occurring through the immediate-return pathway. In contrast, the immediate-return density *f* ^*R*^(*t*) and hazard *h*^*R*^(*t*) remain relatively unchanged. On the low-activity branch, this redistribution produces a progressively larger early peak in the aggregate return time distribution and hazard, while the canonical component retains a long tail. Therefore, subclinical hyperexcitability manifests as an increase in the number of spikes fired as bursts, more spikes sustained per burst, and some non-bursting neurons now starting to fire in bursts. In neurons capable of physiological bursting, the *burst rate* (number of bursts per unit time, *ρ*^*C*^*ρ*^*R*^*/µ*, see Eq. (33) in Methods) may change depending on the relative changes in *ρ*^*C*^, *ρ*^*R*^, and the mean return time *µ*.

In contrast, the jump from the low-to the high-activity attractor is associated with a markedly shorter canonical return time tail *f* ^*C*^(*t*) and a higher canonical hazard *h*^*C*^(*t*); the changes to *f* ^*R*^(*t*) and *h*^*R*^(*t*) are relatively modest. Therefore, an overt hyperexcitable state is marked by a sharp increase in the burst rate, while the ISI of spikes within a burst remain relatively unchanged. Concurrently, *ρ*^*R*^ increases while *ρ*^*C*^ decreases, leading to a further increase in the number of spikes per burst relative to subclinical hyperexcitability, which in turn showed a similar increase relative to normal physiological activity.

Experimental observations support the distinct signatures predicted for the two pathological regimes. In subclinical hyperexcitability, an increased proportion of spikes delivered in bursts relative to controls were observed in mouse models of neuropathic pain [44] and AD [45]. In AD, both patient-derived cortical neurons [46] and CA1 neurons of APP/PS1 mice [45] exhibited more spikes per burst and longer bursts. Acquired intrinsic bursting in previously solitary-firing neurons was also observed in APP/PS1 mice [45], and chronically epileptic rats [47]. Bimodal ISI distributions are also observed during interictal epileptiform discharges [48].

In overt seizures, intracellular recordings from thalamic reticular neurons of GAERS rats showed a switch from short bursts of 2–6 spikes interictally to prolonged bursts of 5–15 spikes during seizures [49], while the burst rate increased nearly seven-fold [50]. Furthermore, after burst stabilization, increasing extracellular potassium in an epileptiform hippocampal slice preparation increased the burst rate without changing the ISI of intra-burst spikes [51], aligning with our steady state observations.

### Immediate-return pathway provides a control dimension on population activity distinct from activation

We next isolate the effect of each pathway on the firing hazard. An earlier placement of the immediate-return pathway or a higher rate of immediate-return leads to the appearance of a high-activity attractor at relatively lower values of the external drive (Figs. 5(a&c)). Varying the parameters corresponding to the immediate-return pathway manipulates both the early and late portions of the return time distribution and the hazard whereas varying those of the canonical pathway influence only the late portions (Figs. 5(b&c) and S3).

**Fig. 5:**
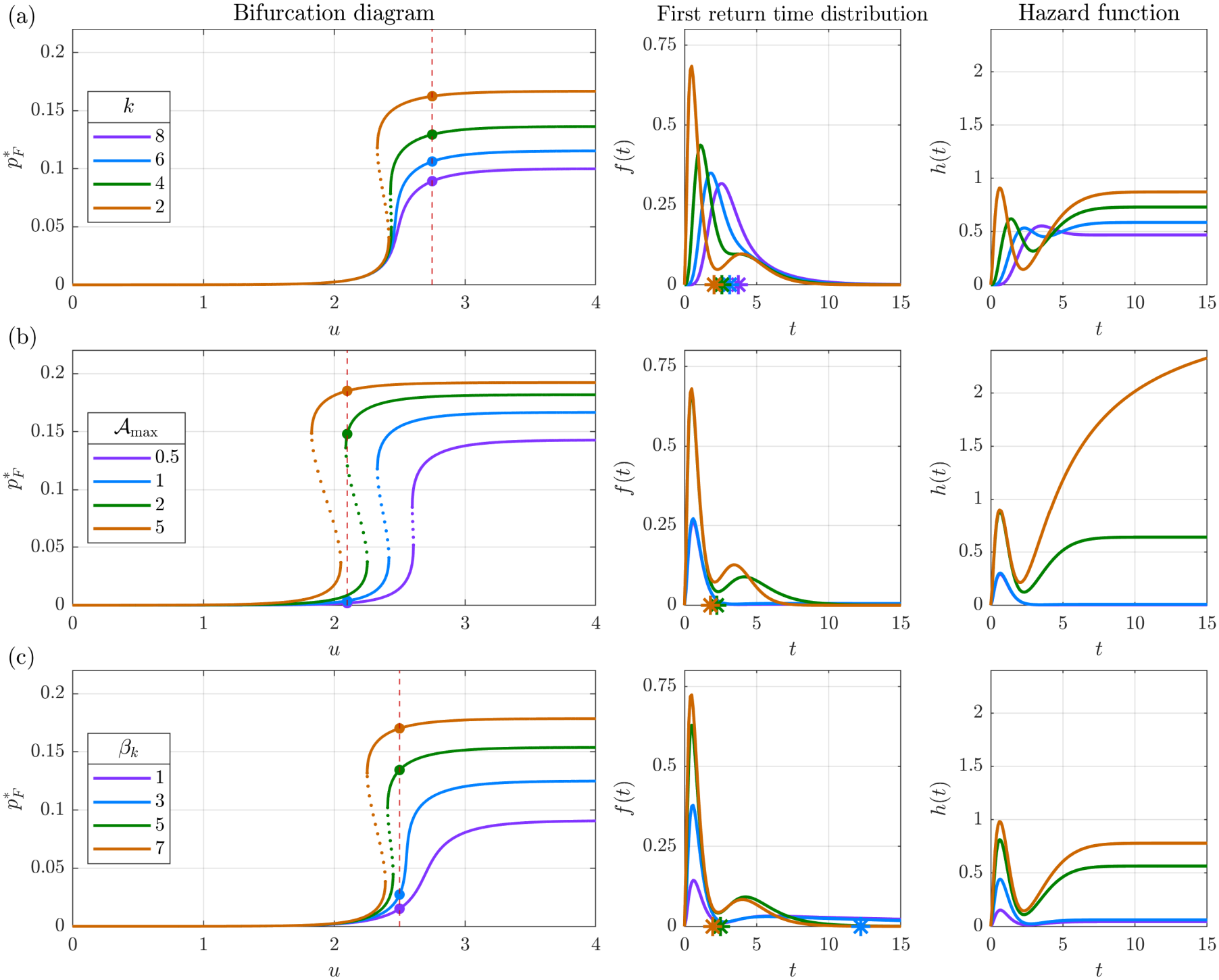
Competing immediate-return and canonical pathways facilitate independent tuning of early and late portions of the return time distribution and the hazard. In each panel, the left subpanels show the 1-D bifurcation diagram for the variation in the steady state firing occupancy 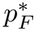 against external drive *u*. Stable branches are indicated in solid lines while the unstable branch is indicated in dotted line. The middle and the right subpanels show the first return time distribution *f* (*t*) (with the corresponding mean return time indicated as an asterisk) and the hazard function *h*(*t*) at select points indicated in the bifurcation diagram. (a) Effect of the location of the immediate-return pathway, with earlier placements leading to higher 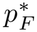. As the immediate-return pathway is moved downstream, it becomes difficult to visually distinguish its contribution from that of the canonical pathway in *f* (*t*) as the time scale separation between the two pathways reduces. However, the aggregate hazard *h*(*t*) still retains its non-monotonicity for the parameters explored. (b) Increasing the supremum of the activation function A_max_ selectively increases the late portion of the return time distribution and the hazard: the early portions remain unaffected among the low-activity attractors (blue and purple curves) as well as among the high-activity attractors (green and brown curves), but differs between the two categories of attractors. (c) Increasing the equivalent parameter for that of the immediate-return pathway *β*_*k*_ modifies both early and late portions of the return time distribution and the hazard even within attractors of the same category. Remaining parameters are explored in Supplementary Figure 3. Panels (a) and (c) further demonstrate that both an earlier placement of the immediate-return pathway or a higher rate of immediate-return lead to the appearance of high-activity attractor at relatively lower values of the external drive. Parameters are set as in Supplementary Material.

Where multiple pathways to firing exist, the first re-entry ends the spike cycle. Therefore, parameters that influence rates downstream of *C*_*k*_ in the canonical pathway — including those of the activation function *A*(*z*^*^) — do not affect neurons that return to *F* directly from *C*_*k*_ along the immediate-return pathway. Therefore, immediate-return retains sole influence over the early portion of the return time distribution (*ρ*^*R*^*f* ^*R*^(*t*)). Since the corresponding contribution to 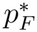 depends only on the mean of this component, *µ*^*R*^ = ⟨*ρ*^*R*^*f* ^*R*^(*t*)⟩, immediate-return provides an independent control dimension over population activity, distinct from activation. This distinction holds even when a sequestration pathway as illustrated in Fig. 2 is added; see Theorem 2 in Methods for proof.

The early hazard component *w*^*R*^(*t*)*h*^*R*^(*t*), however, depends on *A*(*z*^*^) through its weight, *w*^*R*^(*t*) = *ρ*^*R*^*S*^*R*^(*t*)*/* (*ρ*^*R*^*S*^*R*^(*t*) + *ρ*^*C*^*S*^*C*^(*t*) + *ρ*^*S*^*S*^*S*^(*t*)) . When *t* is small, as is for early returns, almost all neurons taking the canonical and sequestration pathways survive, so *S*^*C*^(*t*) ≈ *S*^*S*^(*t*) ≈ 1 over this period. Therefore, by Theorem 2(i), the influence of *A*(*z*^*^) on the early hazard remains negligible.

This independence also survives when the dynamics are projected to 1-D and solely expressed in terms of the firing occupancy alone as in a Wilson–Cowan-type formulation:

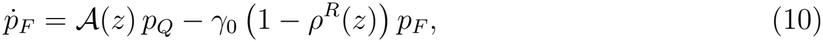

with the quiescent occupancy fixed by conservation,

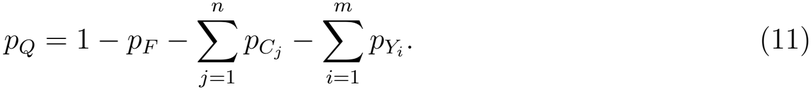

At steady state, the intermediate occupancies 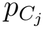 and 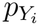 can be expressed in terms of 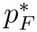 through the renewal-reward theorem, and so, Eqs. (10) and (11) constitute a one-dimensional reduction of the dynamics in *p*_*F*_ . See Theorem refthm:nonlinear-decay in Methods for the derivation.

In Eq. (10), the first term *A*(*z*)*p*_*Q*_ denotes *recruitment* : quiescent neurons are drawn into firing at rate *A*(*z*). The second term *γ*_0_ (1 − *ρ*^*R*^(*z*)) *p*_*F*_ denotes *recurrent firing* : immediatereturn pathway returns a fraction *ρ*^*R*^(*z*) of departing neurons directly from *C*_*k*_ to *F* . Upon projection to 1-D, the excursion *F* → · · · → *C*_*k*_ → *F* collapses to *F* → *F*, and the net decay is reduced accordingly. Thus, the mechanistic content of the immediate-return pathway manifests outside the activation function as effective nonlinear decay.

Substituting for *ρ*^*R*^(*z*) in terms of model parameters, the effective decay is,

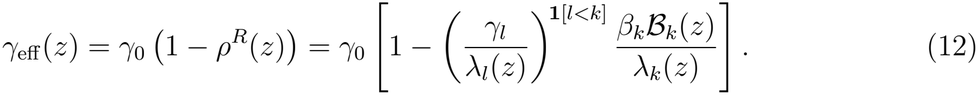

When sequestration lies upstream of immediate-return, the survival factor *γ*_*l*_*/λ*_*l*_(*z*) reduces *ρ*^*R*^ and promotes net decay — neurons diverted onto the sequestration pathway never reach *C*_*k*_ and cannot engage in recurrent firing. Sequestration also opposes firing by depleting the quiescent pool available for recruitment as in Eq. (11), and this second effect operates irrespective of branch order, including when sequestration lies downstream of recurrence.

While such quiescent pool modification is possible through a refractory correction in existing Wilson–Cowan type frameworks, decay is ordinarily a fixed linear relaxation whose role is population bookkeeping: activity not maintained through recruitment leaves the active pool at a constant rate. Our competing hazards formulation mechanistically derives a nonlinear decay and therefore constitutes a generalization of such frameworks.

### Sequestration enables selective suppression of pathological activity

Having explored how competing pathways influence pathological hyperexcitability, we now turn to its selective pharmacological suppression. Use-dependent sodium-channel blockers such as phenytoin and carbamazepine preferentially bind to inactivated sodium channels, delaying recovery and suppressing rapid repetitive firing while largely sparing physiological activity [29, 30]. We represent this action through a sequestration pathway that departs from *C*_*l*_, traverses *m* non-firing, non-recruitable stages, and terminates at *Q* (Fig. 2). Since these drugs exhibit a greater affinity to channels in neurons that have recently fired [31], we place sequestration upstream of immediate-return (*l* = 1, *k* = 2), with *m* = 30 stages and equal exit rates *α*_*i*_ = *γ*. Drug binding is represented by a constant sequestration rate *κ*_*l*_, as binding rate varies linearly with drug concentration [52]. Direct modulation of this rate by ongoing synaptic input is omitted, so *Y*_*l*_(*z*) = 1.

The two-parameter bifurcation diagram (Fig. 6(a)) shows how sequestration (drug action) counteracts an enduring propensity for immediate-return (pathology). Starting from a low-activity attractor representative of physiological bursting (*β*_*k*_ = 2), increasing *β*_*k*_ introduces bistability with the appearance of a co-existing high-activity attractor corresponding to pathological bursting (Fig. 6(c)). Further increase in *β*_*k*_ destroys this bistability, with only the high-activity attractor surviving (Fig. 6(d)). Holding pathology fixed at *β*_*k*_ = 8, increasing intervention through *κ*_*l*_ reverses this progression: a low-activity attractor reappears, followed by the disappearance of the high-activity attractor (Fig. 6(h–e)). Between the untreated high-activity state (Fig. 6(h)) and the treated low-activity state (Fig. 6(e)), firing occupancy decreases by an order of magnitude, from 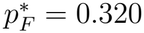 to 0.030.

**Fig. 6:**
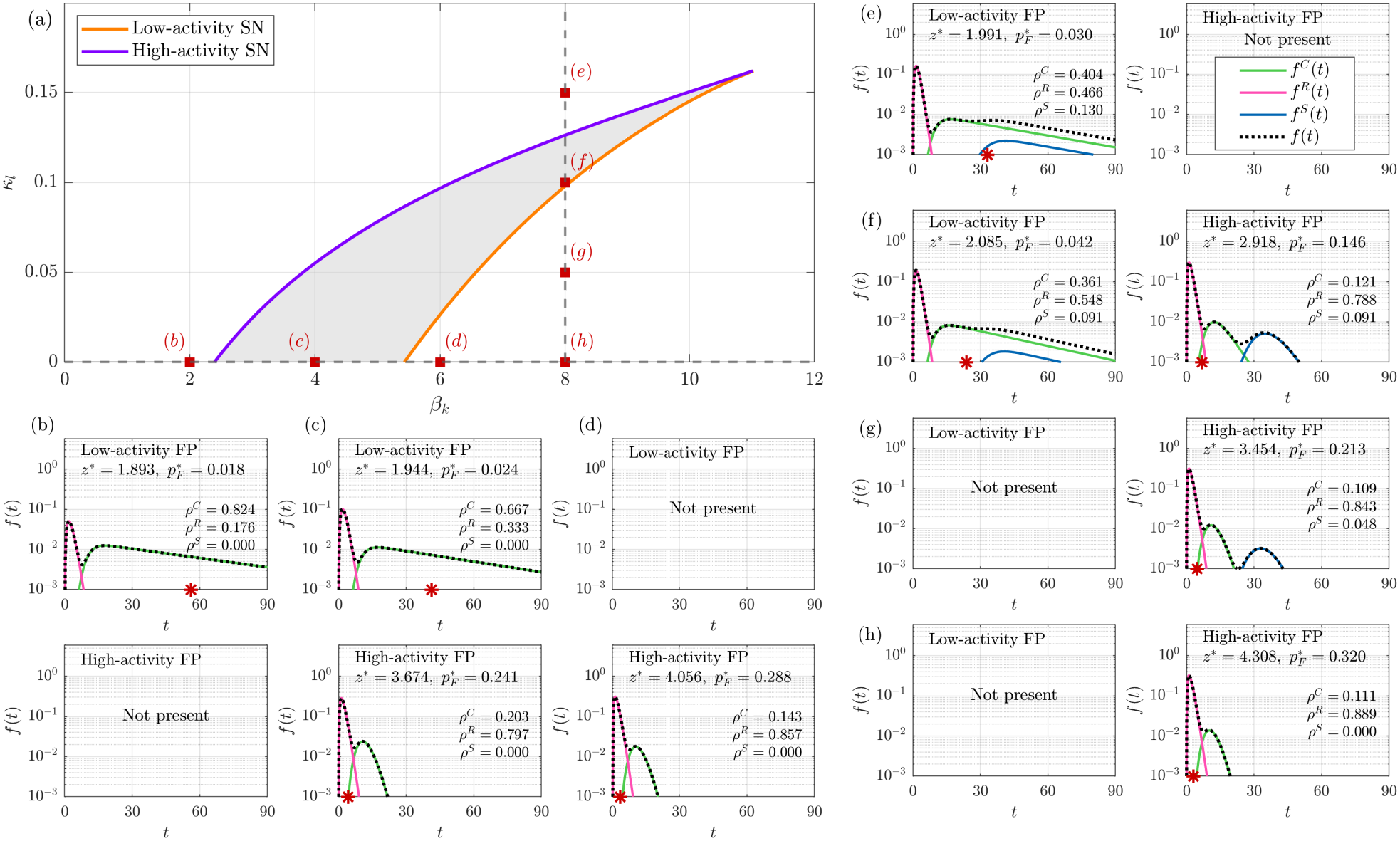
Pathological hyperexcitability hastens the return time distribution and sequestration counteracts it by introducing a late-return component. (a) Two-parameter bifurcation diagram over the rate of immediate-return *β*_*k*_, indicating the level of pathology, and rate of sequestration *κ*_*l*_, indicating the level of intervention, with all other parameters held fixed. Ongoing drive is considered to not influence drug binding, and so Y_*l*_(*z*) = 1. Immediate-return branches from *k* = 2 and sequestration from *l* = 1, and the sequestration pathway consists of *m* = 30 stages with equal exit rates *α*_*i*_ ≡ *α* = *γ*. The loci of saddle-node bifurcations where the high-activity and low-activity attractors lose stability are shown in purple and orange, respectively, and enclose the shaded bistable region. Panels (b to h) illustrate the first return time distribution (dotted line) and its pathway-specific contributions (solid lines), with superscripts *C, R*, and *S* denoting canonical, immediate-return, and sequestration pathways, respectively. Starting from physiological bursting, represented by a low-activity attractor with both canonical and immediate-return components (Panel b), increasing the propensity to take the immediate-return pathway first leads to bistability (c) and subsequently to a monostable pathological high-activity attractor (d and h). This shift is accompanied by a substantial decrease in the mean return time, marked as red asterisk in each panel. Starting from a pathological high-activity attractor (h), increasing the sequestration rate introduces a late-return component, increasing the mean return time (g). Increasing sequestration further brings back bistability (f), and eventually a monostable low-activity attractor representative of physiological bursting (e). During high-activity, the canonical return is sped up and so the canonical and sequestration components can be resolved into separate modes based on the return time distribution alone. However, during low-activity, the canonical return is much slower and therefore the pooled distribution of canonical and sequestration pathways cannot be resolved without information from an additional variable. Parameters are set as in Supplementary Material.

This suppression of pathological bursting by sequestration is expressed through two observables. First, sequestration introduces a much slower component into the return time distribution (Fig. 6(g)). Since the total mass remains conserved, the proportion of neurons sequestered are not available for firing via immediate-return. This reduces *ρ*^*R*^ (Theorem 4 in Methods, illustrated in Supplementary Fig. S4) and therefore the number of spikes sustained per burst. Second, it reduces the burst rate, *ρ*^*R*^(*ρ*^*C*^ + *ρ*^*S*^)*/µ* (see Eq. (33) in Methods), where increasing *ρ*^*S*^ increases the mean return time *µ*.

The resulting drug-treated low-activity state (Fig. 6(e)) therefore resembles physiological bursting but differs from the state before the acquisition of pathology (Fig. 6(b)). In particular, *β*_*k*_ remains elevated during the former, and continued drug exposure offsets the altered balance among pathways and counteracts the expression of pathology. Sequestration thus supports both maintenance and acute seizure termination. As pathology increases, maintenance requires a dose sufficient to sustain the low-activity attractor, whereas, when initiated in high-activity attractor, acute termination requires a dose sufficient to annihilate it.

The apparent sparing of physiological bursting while suppressing pathological bursting by use-dependent sodium channel blockers is explained by how sequestration interacts with the existing return pathways. During high activity, canonical returns are accelerated and so, the late-return component from sequestration appears distinct (Fig. 6(f,g)). During low activity, this component overlaps with the slower canonical contribution and does not form a separate mode that can be distinguished based on interspike intervals alone (Fig. 7(a,b)). Consequently, despite ongoing drug action, the treated and untreated low-activity return time distributions remain broadly similar, giving the impression of normal physiological activity being spared.

**Fig. 7:**
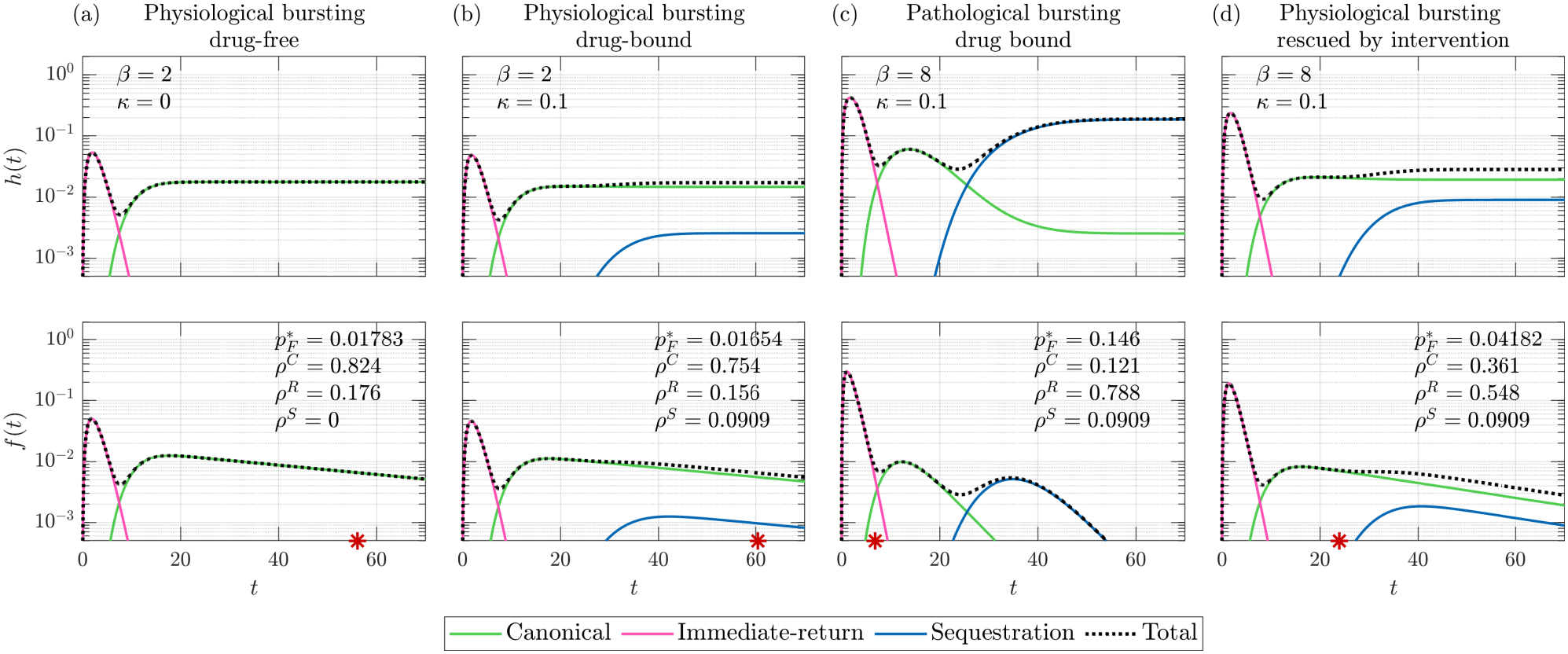
Source of apparent selective pharmacological suppression of hyperexcitability. (a) Drug-free physiological bursting, (b) Drug-bound physiological bursting, (c) Drug-bound pathological bursting, high-activity attractor, and (d) Physiological bursting, where low-activity attractor is rescued by drug in the presence of pathology. For each column, the top row shows the firing hazard and its survival-weighted components while the bottom row shows the return time distribution and its components. The aggregate return time distribution of drug-free (a) and drug-bound (b) physiological bursting remain relatively similar, giving the impression of normal physiological function spared by use-dependent sodium channel blockers such as phenytoin and carbamazepine. On the other hand, the aggregate return time distribution of pathological bursting (c) is quite different from the normal physiological function rescued by the drug (d). Furthermore, during high-activity pathological bursting (c) representative of ictal state, for late returns (*t* ≳ 27), the hazard of firing through the sequestration pathway (as a result of drug binding) exceeds that of the canonical pathway (unbound neurons). In contrast, the sequestration hazard remains below that of the canonical hazard during normal physiological low-activity attractor (b) as well as low-activity attractor rescued by the drug (d). Therefore, most of the late returns are attributable to drug binding during the ictal state but not during the normal or interictal state, despite the same drug concentration. This difference is offered as an experimentally-testable prediction to validate our framework. Parameters are set as in Supplementary Material.

### Competing pathways enable heterogeneity in response in neurons sharing a type and drive

We next illustrate how competing pathways enable heterogeneity in response in neurons of the same population subject to a common drive. Experimental studies indicate that seizures are underpinned by heterogeneous firing patterns among neighboring cells and the changes in the firing rate are not attributable to a homogeneous increase or decrease in the number of active neurons within the recorded population. In human focal seizures, at onset, within a small patch of neocortex, some putative pyramidal cells increased their firing rate while others decreased it. This divergence was observed within a single cell class, and therefore not attributed to interleaved spiking of pyramidal cells and interneurons [33]. Further, in several of the seizures recorded, the firing rate increase during seizure onset was accompanied by a decrease in the number of active neurons, leading to an increase in the Fano factor [33]. Similar heterogeneous shifts in firing rates were observed in excitatory neurons during low-voltage fast activity in human mesial temporal lobe epilepsy [53]. In rodent absence seizures, both thalamic and cortical neurons were observed to fall into one of four distinct patterns: a sustained increase in firing rate over the course of the seizure, a sustained decrease, an onset peak, or no change [34].

Competing pathways provide a mechanism for this divergence. When both immediate-return and sequestration pathways are present within the same population, different neurons sample different pathways on different occasions. To demonstrate this, we simulated 2000 excitatory neurons using the Gillespie method, with quenched parameter heterogeneity, as specified in the Supplementary Material. Entry into immediate-return and sequestration pathways depended on ongoing drive, with thresholds varying between neurons. The common external drive was held at *u* = 1.5 until *t* = 50, briefly raised to *u* = 11.5, and maintained at *u* = 6.5 from *t* = 51 onward.

The increased drive produces a marked increase in population firing occupancy, with an initial peak followed by sustained elevated activity (Fig. 8(a,b)). This aggregate increase conceals divergent neuronal responses. The fraction of uniquely active neurons rises sharply and remains above baseline (Fig. 8(d)), even though some neurons reduce their firing. Relative to each neuron’s mean firing rate over the baseline interval *t* ∈ [10, 50), a majority increase their firing rate by more than a third, while a substantial minority decrease it by a third; the remaining neurons stay within this range (Fig. 8(e)). These differences persist after the drive has settled to its new constant value, demonstrating sustained response heterogeneity under a shared input.

**Fig. 8:**
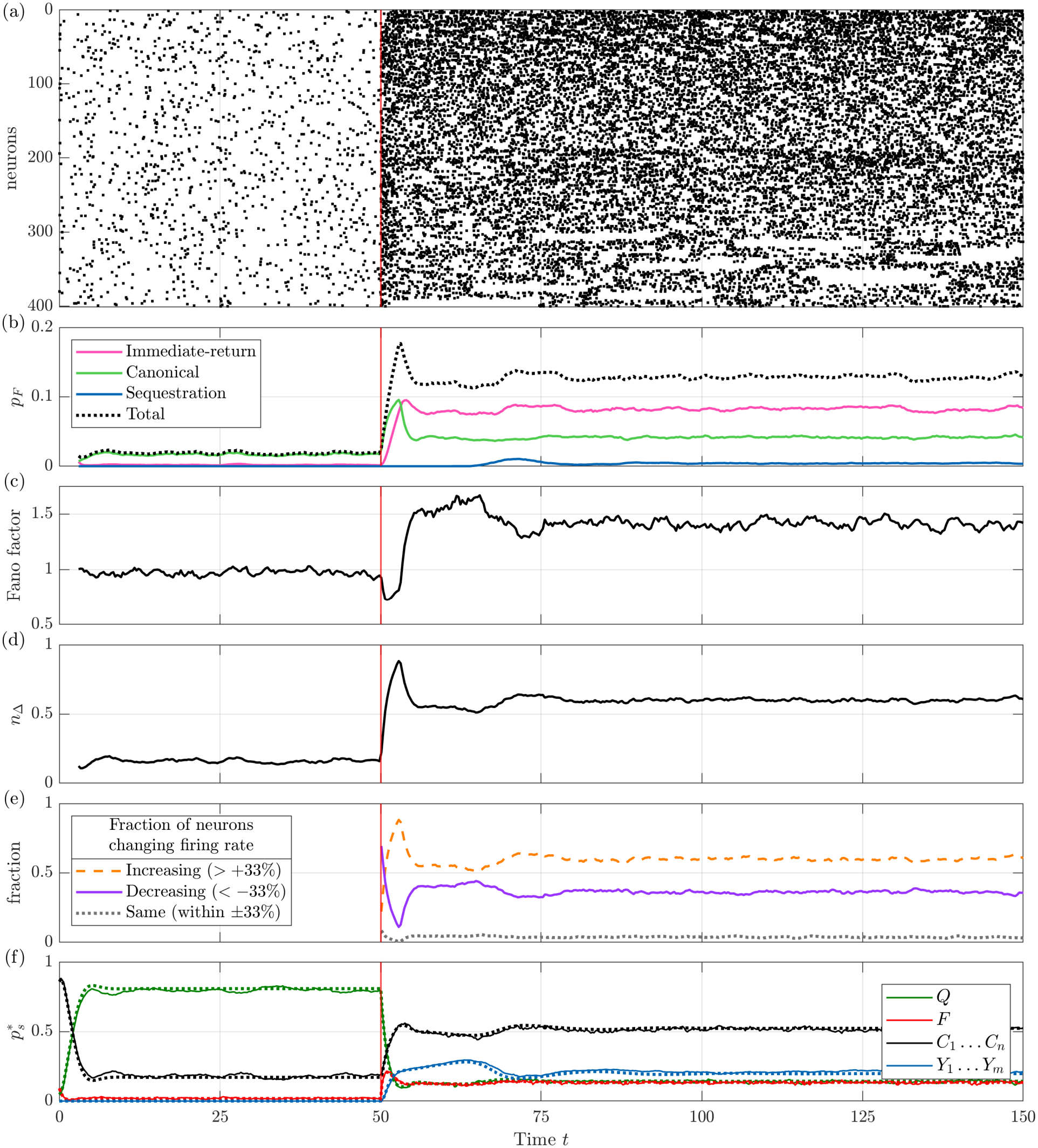
Under a common external drive, competing post-spike pathways enable some neurons within a population to increase their firing rate and some others to decrease it. A population of 2000 excitatory neurons were simulated using the Gillespie method with quenched heterogeneous parameters as described in the Supplementary Material. The external drive was held at *u* = 1.5 from *t* = 0 to *t* = 50. Between *t* = 50 and *t* = 51, the drive was sharply raised to *u* = 11.5. After *t* = 51, the drive was held constant at *u* = 6.5. (a) Raster plot showing the firing activity of 400 neurons chosen at random from the population. (b) Firing occupancy *p*_*F*_ resolved by the pathway taken to firing. (c) Fano factor. (d) Fraction of uniquely active neurons *n*_Δ_ for Δ = 3. (e) Fraction of neurons changing their firing rate relative to their baseline. For each neuron, the baseline firing rate is the mean firing rate calculated over the interval *t* ∈ [10, 50). (f) Occupancies of various states over time, with values from the Gillespie simulation plotted as solid lines and those from the homogeneous mean field equations (Eq. (25)) plotted as dotted lines. The two agree as the dispersion of quenched heterogeneous parameter values is small relative to their mean. The occupancies of canonical intermediate states *C*_1_ to *C*_*n*_, and those of sequestration states *Y*_1_ to *Y*_*m*_ are summed and plotted together for clarity.

Resolving firing occupancy by return pathway shows that immediate-return accounts for most of the increase in firing, while sequestration contributes little to ongoing firing (Fig. 8(b)). Nevertheless, a noticeable fraction of neurons occupy sequestration stages after the input increases (Fig. 8(f)), and remain unavailable for firing. This contrast in response based on the pathway taken leads to the dispersion of neuronal spike counts across the ensemble. The sequestration and immediate-return pathways have return times that differ by an order of magnitude. Therefore, as neurons are increasingly diverted onto these pathways with rise in input, the between-pathway variability increases while the mean return time decreases, leading to an increase in the Fano factor (Fig. 8(c)). Thus, competing pathways introduce a previously unrep-resented form of population organization where neurons of the same type and experiencing a similar drive are separated by which pathway each one takes back to firing.

Rasters simulated from our model (Fig. 8(a)) differ from those obtained experimentally [33] in three aspects, each associated with a modeling assumption: first, successive post-spike cycles retain no explicit memory of the pathway previously taken. Quenched heterogeneity gives neurons persistent pathway preferences, but individual neurons can still take different routes across cycles. Consequently, prolonged silent epochs associated with sequestration occur at different times across the simulated population (Fig. 8(a)). More persistent cell-specific response patterns could be enabled by introducing history-dependent pathway engagement. Next, temporal alignment of responses of cells across the population could be explored by relaxing the weak scaling assumption in Eq. (22). Finally, including interacting excitatory and inhibitory populations would allow these pathway dynamics to be examined alongside inhibition.

## Discussion

Whether a neuron will fire again is fundamentally a different question from whether a quiescent neuron will activate. Therefore, we propose that hyperexcitability is a property of how neurons return to firing from different points in the spike cycle, and not just from quiescence. This temporal structure is lost when population activity is summarized by mean firing rate and interspike interval variability (ISI CV), both commonly used in experimental and modeling studies of hyperexcitability. In the adjacent literature on population spiking, models already retain this structure by tracking the full return time distribution [25, 54].

The firing hazard advances this analysis in two respects. First, a return time distribution can remain unimodal despite comprising distinct return components, whose contributions may become more visually apparent in the corresponding hazard despite a one-to-one correspondence between the two quantities (Fig. 5(a)). Second, the hazard directly describes the instantaneous propensity to fire conditional on not having fired again, which is essentially an age-resolved quantitative description of excitability. We therefore advocate incorporating this description into experimental studies of hyperexcitability.

Doing so would provide a principled basis for understanding how an increased propensity to fire can reflect different alterations across the post-spike cycle, each with different susceptibilities to intervention. At the level of single neurons, a medication may fail to suppress hyperexcitability if it leaves the pathological hazard component largely intact. For instance, prolonged exposure to benzodiazepines can lead to GABA_A_ receptor internalization and reduced responsiveness to the medication [55]. Longitudinal monitoring of the firing hazard can track how the temporal structure of firing propensity changes over the course of treatment, identifying singleneuron signatures of refractoriness and informing the choice of alternatives. The same analysis could support re-purposing anti-seizure medications for hyperexcitability in Alzheimer’s disease and other dementias [56, 57], characterizing not only the suppression of pathological firing but also how interventions alter physiological firing, and whether such alterations are undesirable.

Existing experimental analyses commonly reconstruct the return time distribution from ISIs alone, with some explicitly calculating firing hazards [11]. Recently, Otomo et al. related the ISI distribution to the distribution of the preceding membrane voltage minimum (*V*_min_) in dopamine neurons [32]. They revealed that in pacemaking neurons, both the ISI and *V*_min_ distributions are unimodal, but either or both become bimodal in bursting neurons and quantified this relationship in terms of the CV and the bimodality coefficient.

Here, we take a step further and formally establish non-monotonicity of the firing hazard and resolve it into components distinguished by a physiological measurement other than the ISI itself. This allows the validity of the physiological classification to be tested against the return dynamics. For the hazard components based on *V*_min_ classifications, the associated class weights vary with elapsed time when the components have distinct survival functions (Fig. 4(c)), but remain constant when those functions are identical. Presence of this dependence tests whether the physiological classification distinguishes return dynamics beyond chance. By comparison, the pathway probabilities weighting an ISI mixture remain constant regardless of whether the components are distinct, and supply no equivalent diagnostic (Fig. 4(b)). For the dopamine neuron data [32], 22 out of 23 *V*_min_ two-class representations yielded distinct components based on this test. We provide open-source code so that this analysis can be repeated under other conditions, including comparisons of drug-bound and drug-free neurons to reveal the cause-specific perturbation of excitability in neurons.

We then represent these components as competing hazards and construct a mathematical framework to examine the associated consequences for hyperexcitability. While existing models of physiological [58, 59] and pathological [60, 61] dynamics incorporate a hazard function in deriving the forward dynamics, none make the hazard the explicit object of study. Through a simple renewal framework restricted to steady state, we connect the first return time distribution to the ensemble firing occupancy, bridging microscopic return dynamics to a macroscopic observable of hyperexcitability. Formulating the problem in terms of return times rather than forward dynamics allowed us to derive closed-form analytical results for entire classes of models.

We distinguish two processes that are often confounded in models of hyperexcitability: recruitment of a neuron that has been quiescent for a while versus the recurrent firing of a recently active neuron. We show that this distinction survives projection onto a 1D Wilson-Cowan-type formulation, appearing outside the activation function as a nonlinear decay. Hyperexcitability reduces this decay by routing recently active neurons toward recurrent firing. In an earlier work, we showed that a second-order decay term, imposed by hand in a Wilson–Cowan-style model, could account for seizures that resist benzodiazepines yet terminate with phenytoin or carbamazepine [62]. This emphasizes the importance of nonlinear decay, and identifies a new locus of hyperexcitability that could be incorporated in existing formulations.

Resolving this distinction has also enabled us to distinguish subclinical from overt hyper-excitability: relative to baseline physiology, subclinical hyperexcitability leads to more spikes delivered in bursts [44, 45], including the conversion of solitary-firing neurons into bursters [47]. Overt hyperexcitability, on the other hand, substantially increases the bursts fired per unit time [49, 50].

Pathway competition also enables selective pharmacological suppression of pathological hyperexcitability when a fraction of neurons are diverted onto prolonged inactivity, for instance, by use-dependent sodium channel blockers such as phenytoin and carbamazepine. Levetiracetam also enables selective pharmacological suppression [63]. Rather than influencing the return time distribution, we posit that it affects the self-consistent feedback within the population by gating the release of the excitatory neurotransmitter glutamate and leave further analyses to future work.

Finally, experiments have shown that seizures are underpinned by heterogeneous firing patterns among neighboring cells within the same population, where some neurons increase their firing rates, while others decrease theirs [33, 34]. Truccolo et al. noted that different groups of neurons appeared to enter and dwell in different states with different time constants [33]. Competing pathways provide a concrete realization of this account: neurons sharing a type and drive differ in their engagement of routes with distinct return times, allowing some to sustain recurrent firing while others enter prolonged inactivity. While heterogeneity emerging from network effects both within a neuronal population and between populations of different cell-types have been extensively studied [64, 65], one arising from how a neuron returns to firing remains unexplored. Introducing post-spike pathway heterogeneity therefore distinguishes our contribution from comparable frameworks.

In terms of limitations, the analytical results rely on a renewal description at steady state. This makes the relationship between return dynamics and population activity tractable, but does not capture adaptation and other forms of accumulated cellular history that could modify both successive pathway selection and return times. We also omit spatial organization within the population.

The pharmacological interventions are represented phenomenologically rather than through explicit channel-binding kinetics or drug concentration dynamics. The maintenance and termination thresholds therefore describe intervention requirements within the model, rather than clinical doses. The pharmacological interpretation also remains to be tested by estimating pathway-specific return dynamics in pathological populations and measuring how interventions alter them.

Therefore, we propose a falsifiable prediction that concerns a change in which pathway dominates late returns, despite unchanged drug exposure. At a fixed sodium-channel blocker concentration, during high-activity (ictal state), sequestration should make a greater survival-weighted contribution to the late firing hazard than the canonical pathway (Fig. 7(c)). During low-activity (interictal state) at the same concentration, this ordering should reverse (Fig. 7(d)). Testing this prediction would require recording spike times together with drug-binding history or a suitable proxy like spike rise rate [66], allowing return intervals to be classified according to whether they involved sequestration. This test applies where interictal firing is not substantially suppressed: at sufficiently high drug concentrations, sequestration dominates returns at all activity levels, silencing activity, eliminating the proposed distinction.

In conclusion, here we distinguish excitability arising from more neurons firing versus that from the same neurons firing more often based on post-spike pathway competition. We demonstrate how the hazard of firing quantifies this distinction in whole cell recordings of dopamine neurons. We then use this distinction to identify signatures of overt and subclinical hyperexcitability, their pharmacological suppression, and the associated heterogeneity in within-population response, consistent with experimental findings across epilepsy, Alzheimer’s disease, and neuropathic pain.

## Methods

### Experimental data analysis

For each cell, we tested whether the ISI hazard is monotone, divided the intervals into distinct classes by the trough voltage *V*_min_ preceding each spike, and decomposed the pooled hazard into class-specific contributions. Cells were then grouped by these results and compared.

### Data and spike detection

We re-analyzed *in vivo* whole-cell recordings of spontaneously active midbrain dopamine neurons from Otomo et al. [32]. These were current clamp recordings at zero holding current from tyrosine hydroxylase-identified neurons in isoflurane-anesthetized adult C57BL/6 mice. Spikes were detected using the identical detection method published in Otomo et al. [32]. Interspike intervals (ISIs) were calculated as the differences in time between successive spike peaks. *V*_min_ was the minimum of the filtered trace between two successive spike peaks. Cells with fewer than 80 intervals were excluded from our analysis, leaving 97 of 110 original recordings.

### Discharge variability

To measure the global variability of spiking activity, for each cell, we calculate the coefficient of variation of the ISIs, CV = SD*/*mean. However, CV can be inflated when firing rates are modulated slowly. Thus, we also calculate a measure of local variability which is more robust to these modulations [41],

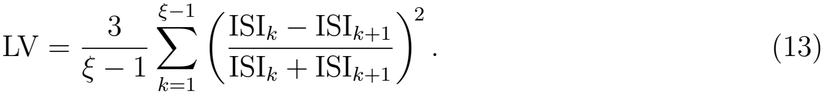

where, *ξ* is the number of intervals in a cell. LV is zero for regular firing, i.e., all ISIs are equal, and one for a homogeneous Poisson process.

### Test for a monotone hazard

To determine whether the ISI hazard was non-monotone, we utilized an isotonic projection test [40, 67]. This test finds the best-fit non-decreasing hazard and assesses whether departures from this fit are explainable by noise. The method was applied to bottom 95% of the ISIs for each cell, rescaling each ISI by the 95th-percentile value, *c*. Let *F*_*ξ*_ be the empirical distribution function of the ISIs. The empirical cumulative hazard is given by *H*_*ξ*_(*t*) = − log{1 − *F*_*ξ*_(*t*^−^)}. Any nondecreasing hazard yields a convex cumulative hazard, as the hazard is its derivative. Thus, we project *H*_*ξ*_ onto the space of convex functions by finding its greatest convex minorant, GCM(*H*_*ξ*_). This gives a distribution function with a nondecreasing hazard that best fits the data, 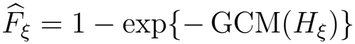. The test statistic

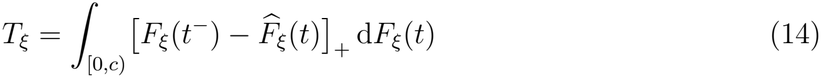

measures how much the data differs from the nondecreasing fit. It is zero when the hazard is nondecreasing and gains positive contributions over subintervals of [0, *c*) where the empirical hazard decreases. To determine whether *T*_*ξ*_ is larger than sampling noise alone would produce, we compared it with its null distribution under a nondecreasing hazard, estimated with the smoothed bootstrap of Groeneboom and Jongbloed [67]. Here, a smooth nondecreasing hazard was fit to the data and rescaled so that its cumulative hazard at *c* equals that of the data, *H*_*ξ*_(*c*). From this null hazard we drew *B* = 1000 synthetic sets of *ξ* ISIs, the same number as in the data, and computed the statistic 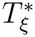 for each. The Monte Carlo *p*-value, 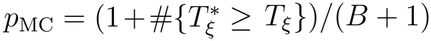, is the fraction of synthetic sets that depart from monotonicity at least as much as the data, counting the data itself so that *p*_MC_ > 0. Cells were classified as non-monotone when *p*_MC_ < 0.05.

Crucially, this method is robust to noise-driven fluctuations, so small decreases in the empirical hazard do not affect the assessment of monotonicity. This test was applied independently of the kernel estimates used for plotting and decomposition of the hazard.

### Classification of intervals by afterhyperpolarization depth

To determine whether a neuron followed distinct post-spike pathways, we divided each cell’s intervals into classes according to the associated *V*_min_. The *V*_min_ distribution was first smoothed with a Gaussian kernel density estimate with bandwidth 1.06 SD *ξ*^−1*/*5^ [68]. Peaks of the density were determined to correspond to classes if they had a peak prominence of at least 10% of its height. For each cell, at most five modes were retained. Intervals were then grouped into classes by segmenting the distribution at the minimum of the density between selected peaks. Classes that contained fewer than 3% of the total intervals were merged into the class with the closest mean *V*_min_.

When the density method yielded only a single class but the empirical hazard was found to be non-monotone, we applied a shoulder criterion instead. We centered a Gaussian with standard deviation 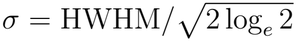 at the maximum of the density, where HWHM denotes half-width at half-maximum. Values lying more than 1.5*σ* from the maximum of the density were counted on either side. If at least five such values were found, we employed a two-sided exact binomial test to check whether they were distributed evenly on both sides of the Gaussian. For a p-value less than 0.01, the distribution was split into two classes at 1.5*σ* on the side with more values. Of the 23 non-monotone cells with two *V*_min_ classes, 12 were identified by the density criterion and 11 by the shoulder criterion.

### Survival, hazard, and class decomposition

Next, we estimated the survival and hazard functions for each cell. At each distinct ISI length observed, we divided the number of ISIs of that length by the number of those that were at least that long (the Nelson–Aalen estimator [69, 70]). We then smoothed the result into a continuous curve with an Epanechnikov kernel [71] of bandwidth 1.35 min(SD, IQR*/*1.349) *ξ*^−1*/*5^. ISIs cannot be negative, so the kernel was reflected about *t* = 0 to avoid underestimating the hazard at short ISIs [72]. To understand how each *V*_min_ class contributes to the hazard, we repeated this process using only the ISIs of each class. All estimates were computed up to the 99.5th percentile of the cell’s ISIs, or until fewer than ten ISIs remained.

Let *ρ*^*j*^ be the fraction of a neuron’s ISIs that belong to the *j*th class, and let *S*^*j*^ and *h*^*j*^ be the class-specific survival and hazard. The total survival is given by the sum over classes,

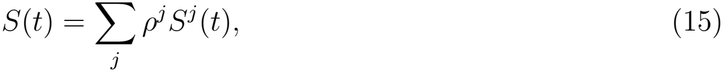

and its hazard decomposes into contributions from each class,

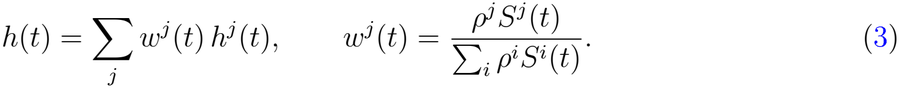

Here *w*^*j*^(*t*) is the fraction of ISIs belonging to class *j* with length at least *t*. It was obtained by dividing the number of ISIs in class *j* by the aggregate number of ISIs at least that long for each observed ISI length. The result was smoothed with the cell’s kernel and bandwidth. Each contribution *w*^*j*^*h*^*j*^ was thus estimated directly.

To test if the classes are truly distinct, we used the fact that if all classes had the same hazard, Σ_*j*_ *ρ*^*j*^*h*^*j*^ would equal *h*. We measured how far Σ_*j*_ *ρ*^*j*^*h*^*j*^ departs from *h* as the root-mean-square relative difference, ignoring times where *h* was below 0.1% of its maximum. We constructed a null by shuffling the class labels 200 times with class sizes fixed, and recomputed it. The decomposition of the hazard was retained if at most four of these shuffles produced a discrepancy as large as the observed.

### Agreement between the *V*_min_ and ISI partitions

We assessed whether the class decomposition from the *V*_min_ distribution correspond directly to those from the ISI length, by dividing each cell’s log_10_(ISI) into classes, using the same method as for *V*_min_ [73]. Each ISI for a neuron was therefore given two class labels, one from its *V*_min_ distribution and one from its log_10_(ISI) distribution. When both methods yielded same number of classes, we paired the class labels so that they maximally overlapped. We measured their agreement with Cohen’s *κ* test [74], where *κ* is 1 when the classifications are identical and 0 when their difference is explainable by noise. The value of *κ* was interpreted with the bands of Landis and Koch [75]. Cells where the two methods produced different numbers of classes were excluded from this comparison. To measure how closely *V*_min_ follows log_10_(ISI) within each cell, we also computed Spearman’s rank correlation between them.

### Statistics

Cells were assigned to the monotone group if the test for a monotone hazard did not reject monotonicity and *V*_min_ had a single class (59 cells), and to the non-monotone group if the test rejected monotonicity, *V*_min_ had two classes, and these classes passed the permutation test of the class decomposition (22 cells). All other cells were excluded from group comparisons. Differences in variability and in Spearman correlation between the two groups were assessed with two-sided Mann–Whitney *U* tests [38] with effect sizes provided by Cliff’s *δ* [39].

### Model and mathematical framework

We define the single-neuron Markov chain and its mean-field population limit, obtain the steady states from the renewal structure of the interspike interval, decompose the interval into pathways, and prove the properties of the hazard used in the Results.

We consider a population of *N* neurons, each occupying an element of the state space

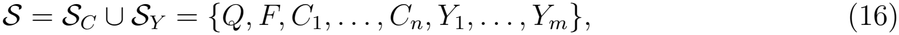

with states: quiescent *Q*, firing *F*, a canonical sequence of post-spike recovery states *C*_1_, …, *C*_*n*_, and a sequestration chain *Y*_1_, …, *Y*_*m*_. The state of neuron *i* at time *t* is 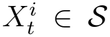, and the joint state 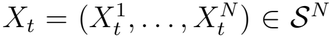 is assumed to evolve as a continuous-time Markov chain (CTMC). The *state occupation probability*

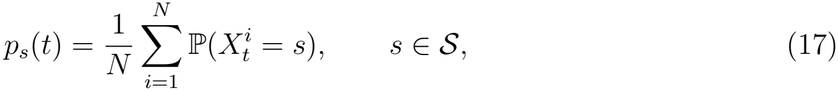

is the probability that a neuron chosen uniformly occupies state *s*, and

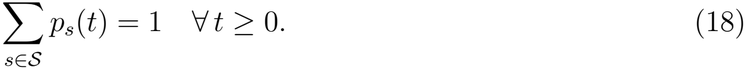

### Transition structure and the generator

A neuron receiving input *z* ∈ ℝ moves from state *s* to *s*^′^ ≠ *s* at the rate 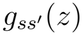. The function 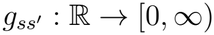 is set to be identical for every neuron in our model, so that neurons’ transition rates differ across the population only through their input. These rates form the single-neuron *generator G*(*z*), a square matrix with rows and columns corresponding to the states with diagonal entries 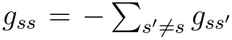. This requirement makes each row sum to zero, satisfying conservation of probability. The time that a neuron stays in state *s*, known as the *sojourn time* in *s*, is distributed exponentially with rate |*g*_*ss*_|, and the neuron jumps to state *s*^′^ with probability 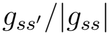. We choose each rate to be either zero for all *z* or positive for all *z*, so the allowed transitions (positive rate) form a fixed directed graph G with vertices corresponding to *S* and edges corresponding to the allowed transitions. We refer to the edge set as the graph’s *topology*.

The graph for the canonical pathway G_*C*_ is simply the cycle *Q* → *F* → *C*_1_ → · · · → *C*_*n*_ → *Q*, with constant recovery rates *γ*_0_, …, *γ*_*n*_ > 0, where *γ*_*i*_ is the rate for *C*_*i*_ → *C*_*i*+1_, identifying *F* and *Q* as *C*_0_ and *C*_*n*+1_, respectively. The input-dependent edge *Q* → *F* is governed by the activation function *A*(*z*) > 0, which is positive, monotonically increasing, and bounded above by *A*_max_. On *S*_*C*_ = {*Q, F, C*_1_, …, *C*_*n*_},

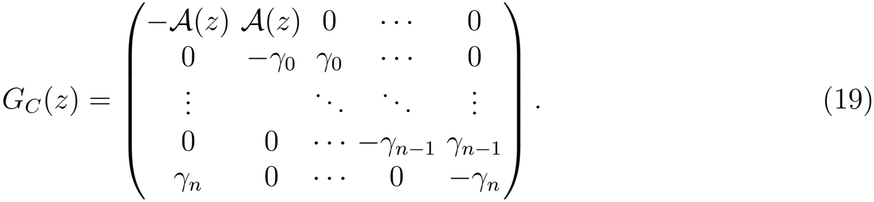

Deviations from this trajectory correspond to *topological perturbations*, which add edges to G_*C*_. Writing **e**_*s*_ for the standard basis vector for the state space indexed by *s*, an *immediate-return* perturbation adds the edge *C*_*k*_ → *F*,

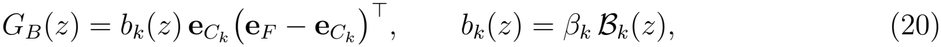

with *β*_*k*_ ≥ 0 and *ℬ*_*k*_ monotonically increasing. *Sequestration* perturbations create branches from *C*_*l*_ through a chain of ordered states, *S*_*Y*_ = {*Y*_1_, …, *Y*_*m*_}, and exits at *Q*. The transition rate from *C*_*l*_ → *Y*_1_ is *y*_*l*_(*z*) = *κ*_*l*_ *Y*_*l*_(*z*), with *κ*_*l*_ ≥ 0 and *Y*_*l*_ monotonically increasing. The rate of exit from *Y*_*i*_ is given by *α*_*i*_ > 0 for *i* = 1, …, *m*. Sequestration then adds

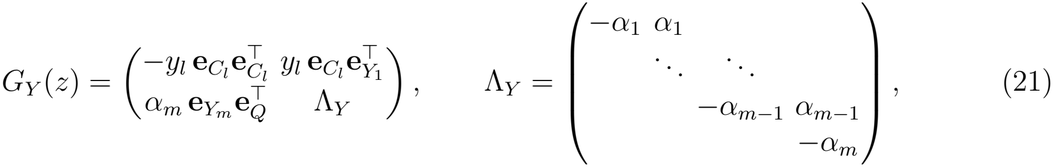

so that 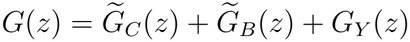, where the tilde denotes padding with zeros to *S*.

### Input and mean-field dynamics

When neuron *j* occupies the firing state *F*, it provides input to neuron *i*, so that

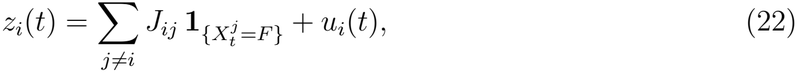

with synaptic weights *J*_*ij*_ ∈ ℝ and external drive *u*_*i*_(*t*). We assume a common drive *u*_*i*_(*t*) = *u*(*t*) and weak scaling, i.e., *J*_*ij*_ = *a*_*ij*_*/N* with *a*_*ij*_ bounded independently of *N* . The scaled weights *a*_*ij*_ are drawn independently with mean *a* and finite variance. A standard average over the quenched disorder replaces *a*_*ij*_ by *a* [76, 77]. The input *z*_*i*_ is then an average over *N* neurons, each scaled by 1*/N* . Thus, as *N* → ∞, *z*_*i*_ concentrates at its mean,

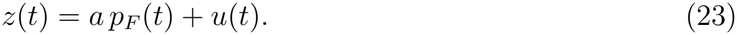

For finite *N*, this neglects fluctuations of *z*_*i*_ about *z* and correlations between neurons [78, 79]. In this mean-field limit, neurons experience identical input, and each evolves as a CTMC with generator *G*(*z*(*t*)) [78, 80, 81]. It follows that the state occupation probabilities **p** = (*p*_*s*_)_*s*∈*S*_ obey the Kolmogorov forward equations [82]

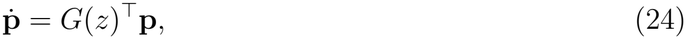

which are generally nonlinear because *z* depends on *p*_*F*_ . With 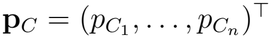 and 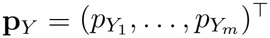, Eq. (24) reads

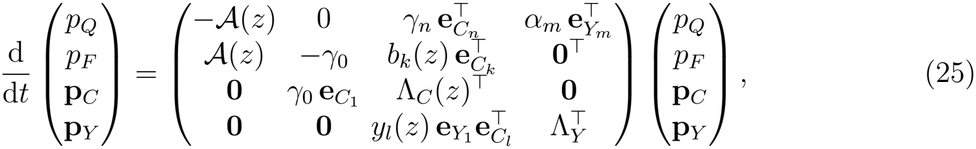

where the recovery-chain sub-generator Λ_*C*_(*z*) is bidiagonal, with diagonal −*λ*_1_, …, −*λ*_*n*_ and superdiagonal *γ*_1_, …, *γ*_*n*−1_. Finally,

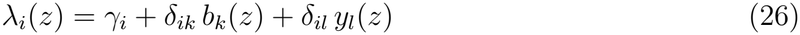

is the total exit rate from *C*_*i*_.

### Renewal structure and steady states

We now consider steady states of the system, **p**(*t*) = **p**^*^, with constant drive *u*(*t*) = *u*. At steady state, the input is constant and given by

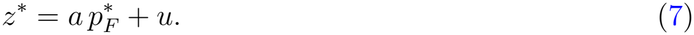

Further, each neuron evolves as a time-homogeneous CTMC with generator *G*(*z*^*^). We note that every state leads to *F* through a directed cycle of G, i.e., a repeating sequence of allowed transitions. We define *T*_*F F*_ to be the time it takes between consecutive returns to the firing state. Its distribution, called the *return time distribution*, has probability density *f* (*z, t*) and finite mean ⟨*T*_*F F*_ (*z*)⟩ = *µ*(*z*), where ⟨·⟩ denotes the expectation over this distribution.

Let *π*_*s*_ be the probability that the neuron enters state *s* during a single return to *F*, and let *τ*_*s*_ = 1*/*|*g*_*ss*_(*z*^*^)| be its mean sojourn time. Thus, *π*_*s*_*τ*_*s*_ is the expected time spent in *s*.

By positive recurrence, the chain has a unique stationary distribution 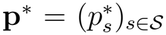 such that *G*(*z*^*^)^⊤^**p**^*^ = **0**. Each component is given by [82]

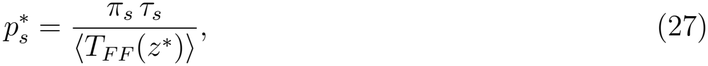

and, since *π*_*F*_ = 1 and *τ*_*F*_ = *γ*^−1^,

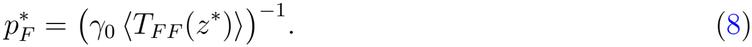

Writing 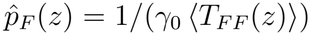, steady states are identified as the fixed points of the self-consistency map 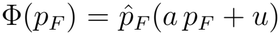. For a given drive *u*, there can be several fixed points, 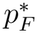. Since each *z*^*^ determines the steady-state firing occupancy, 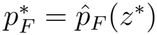, and 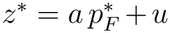, the corresponding drive is

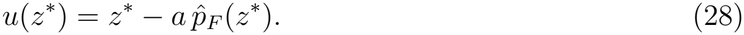

Therefore, the curve (*u*(*z*^*^), 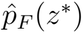 traces every steady state as *z*^*^ is varied.

A perturbation *δp*_*F*_ of the steady-state firing occupancy changes the input by *a δp*_*F*_, which moves the renewal steady state to 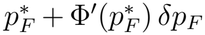. We assume that the remaining occupancies relax quasi-statically to this point, so that, to first order, the perturbation obeys

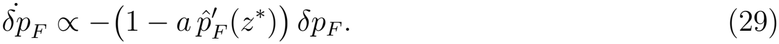

Therefore, under this quasi-static assumption, a steady state is stable if and only if 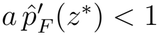, which is equivalent to d*u/*d*z*^*^ > 0. In the full system, d*u/*d*z*^*^ < 0 still implies instability, but oscillatory instabilities are not excluded where d*u/*d*z*^*^ > 0. Since 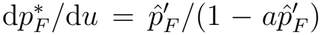, stable states are those that have positive slope along the solution curve in the 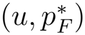 plane wherever 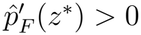.

### Pathway decomposition of the return time

Each cycle begins with a sojourn in *F* distributed exponentially with rate *γ*_0_, Exp(*γ*_0_). There are three possible pathways to return to *F* : canonical (*F* → *C*_1_ → · · · → *C*_*n*_ → *Q* → *F* ), immediate-return (*F* → *C*_1_ → · · · → *C*_*k*_ → *F* ), or sequestration (*F* → *C*_1_ → · · · → *C*_*l*_ → *Y*_1_ → · · · → *Y*_*m*_ → *Q* → *F* ). A neuron exits *C*_*i*_ after an Exp(*λ*_*i*_)-distributed sojourn and moves to the *C*_*i*+1_ (where *C*_*n*+1_ = *Q*), to *F*, or to *Y*_1_ with probabilities *γ*_*i*_*/λ*_*i*_, *δ*_*ik*_*b*_*k*_*/λ*_*k*_, and *δ*_*il*_*y*_*l*_*/λ*_*l*_, respectively. Thus, a cycle reaches *C*_*i*_ with probability 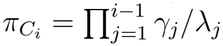. The *pathway probabilities* for canonical, immediate-return, and sequestration are given by 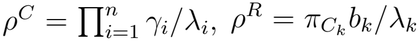, and 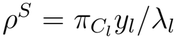, respectively, which must sum to one, as these are the only allowed pathways. These probabilities can all be written explicitly in terms of transition rates,

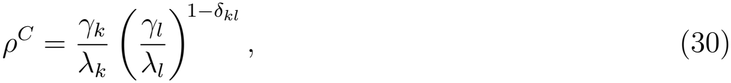

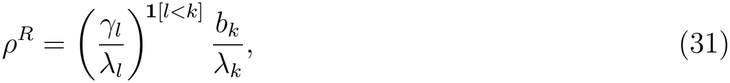

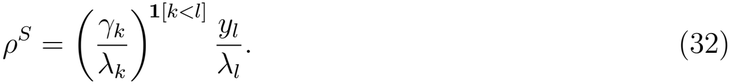

For a positive immediate-return rate (*b*_*k*_ > 0), *ρ*^*R*^ decreases strictly with *y*_*l*_ when *l* ≤ *k*, i.e., sequestration is at or upstream of immediate-return, and does not depend on it when *l* > *k*.

At steady state, each entry into *F* starts the chain anew, because its future depends only on the current state. Successive returns therefore choose their pathways independently, and they occur at mean rate ⟨*T*_*F F*_ ⟩^−1^ [43]. A burst begins when an immediate-return follows a slower canonical or sequestration return, so the *burst rate*, the number of bursts per unit time, is

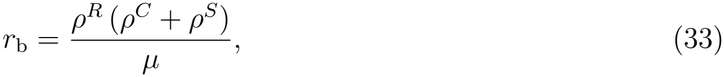

which reduces to *ρ*^*R*^*ρ*^*C*^*/µ* without sequestration. The visit probabilities and sojourn times for state *s* (*π*_*s*_, *τ*_*s*_) (Eq. (27)) are (1, 1*/γ*_0_) for *F*, (*π*_*C*_, 1*/λ*_*i*_) for *C*_*i*_, (*ρ*^*S*^, 1*/α*_*j*_) for *Y*_*j*_, and (*ρ*^*C*^ + *ρ*^*S*^, 1*/A*) for *Q*.

When restricted to a single pathway, the return time to firing is a sum of independent, exponentially distributed sojourns, one for each state visited. Therefore, the *pathway-conditional return time distributions* are given by convolutions,

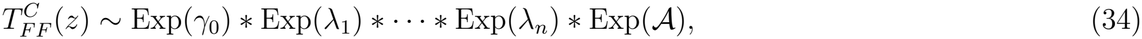

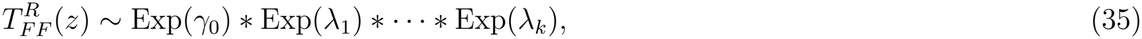

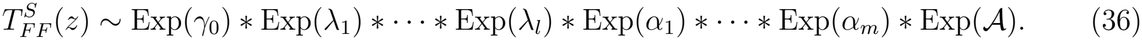

We define the *pathway density f*^*j*^(*z, t*) for *j* ∈ {*C, R, S*} to be the density associated with the distribution of 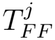. Let 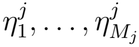 be the exit rates of the states visited along pathway *j*. Let Λ^*j*^ be the *M*_*j*_ ×*M*_*j*_ bidiagonal matrix with diagonal 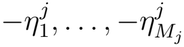 and superdiagonal 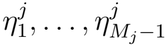. It follows that each *f*^*j*^ is hypoexponential and can be evaluated as 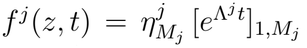, where *A*_*i,j*_ denotes an element of the matrix *A*. The density of *T*_*F F*_ is therefore the mixture

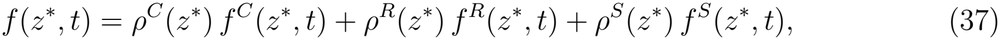

with survival function *S* = Σ_*j*_*ρ*^*j*^*S*^*j*^, where *S*^*j*^ is the survival function of pathway *j*. Its hazard decomposes as *h* = Σ_*j*_ *w*^*j*^*h*^*j*^, where *h*^*j*^ = *f*^*j*^*/S*^*j*^ is the individual pathway hazard with the associated weight *w*^*j*^(*t*) = *ρ*^*j*^*S*^*j*^(*t*)*/S*(*t*).

### Hazard of a single pathway

Each pathway return time in Eqs. (34) to (36) is a sum of independent exponentially distributed sojourn times. The following theorem restates known results for such sums [28].

#### Theorem 1

(Monotone hazard of a single pathway) *Let* 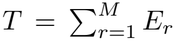 *with independent random times E*_*r*_ ∼ Exp(*η*_*r*_). *Then the hazard h*_*T*_ *is nondecreasing and tends to the smallest exit rate*,

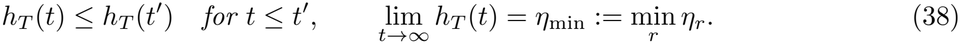

*Consequently, the hazard and survival function are bounded by*

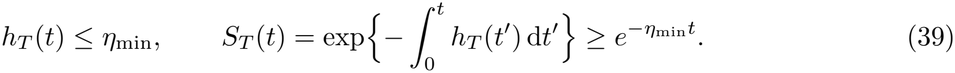

*Proof* The density *f*_*T*_ is log-concave [83] and Eq. (38) immediately follows [28, 43]. Since *h*_*T*_ is nondecreasing, it never exceeds its asymptotic limit, which gives Eq. (39).

For the canonical restriction (*b*_*k*_ = *y*_*l*_ = 0), every cycle follows the same pathway. Hence, the hazard is always nondecreasing, with

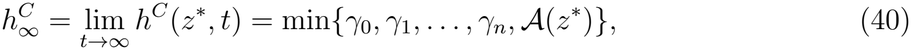

and 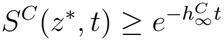. A non-monotone hazard can therefore only exist when there are at least two pathways back to firing.

With immediate-return present and no sequestration (*b*_*k*_ > 0, *y*_*l*_ = 0), the two pathways share every sojourn up to and including that in *C*_*k*_, so 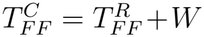 with *W* ≥ 0 independent of 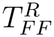. Letting *f* ^*R*^ = 0 and *S*^*R*^ = 1 when their arguments are negative, *h*^*C*^(*t*) = ⟨*f* ^*R*^(*t* − *W* )⟩*/*⟨*S*^*R*^(*t* − *W* )⟩ is a weighted average of *h*^*R*^(*t* − *W* ) ≤ *h*^*R*^(*t*) by Theorem 1. Therefore, *h*^*C*^(*t*) ≤ *h*^*R*^(*t*). Since d(ln *w*^*R*^)*/*d*t* = *h* − *h*^*R*^ = *w*^*C*^(*h*^*C*^ − *h*^*R*^) ≤ 0, the weight *w*^*R*^(*t*) decreases and *w*^*C*^(*t*) = 1 − *w*^*R*^(*t*) increases with *t*.

### Moments of the return time

As expectation is linear, the mean pathway return time is given by the sums of the mean sojourn time in each state,

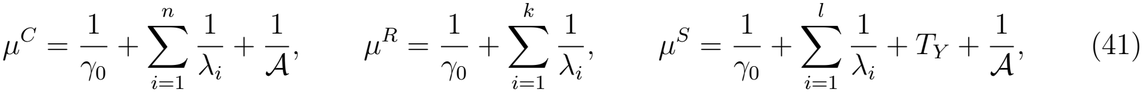

with 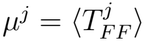 and 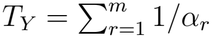 the mean residence time in the sequestration chain. The mean return time is then simply the sum of the mean pathway times, weighted by the pathway probabilities (Eqs. (30) to (32)),

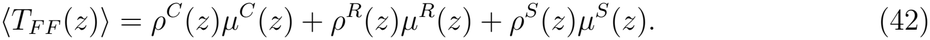

Writing out terms explicitly and collecting them by state,

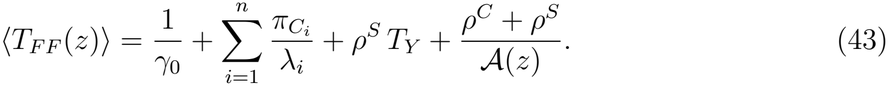

The pathway return time variances 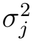 are the sums of the individual state variances. By the law of total variance, 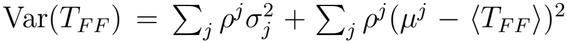 is the sum of within- and between-pathway contributions.

#### Canonical restriction

Without immediate return or sequestration, *ρ*^*C*^ = 1 and *λ*_*i*_ = *γ*_*i*_. Hence, the return time is the sum of independent exponential sojourn times in the states along the canonical pathway. Thus, its mean and variance are the sums of the sojourn means and variances,

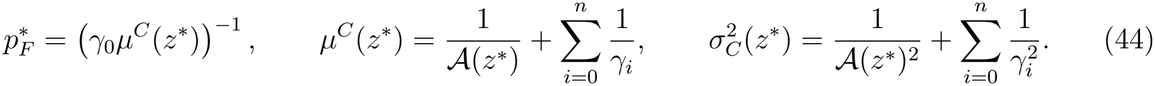

### Independence of the immediate-return pathway from activation

#### Theorem 2

(Independence of the immediate-return pathway from activation) *At fixed z, treat* A = A(*z*) *as a free parameter. Then:*

i. *the pathway probabilities ρ*^*C*^, *ρ*^*R*^, *ρ*^*S*^ *and the immediate-return density f* ^*R*^ *do not depend on A;*
ii. *activation enters the mean return time only through a single term*,

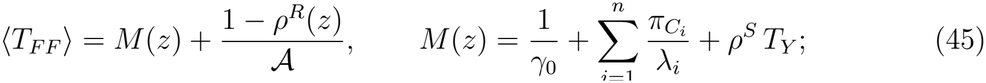
iii. *the sensitivity of the firing occupancy to activation*,

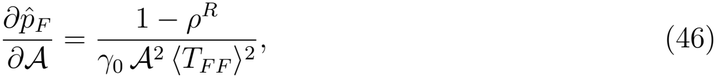

*vanishes as ρ*^*R*^ → 1.

*Proof* (i) *A* is the rate of *Q* → *F* alone and enters none of the exit rates *λ*_*i*_. The pathway probabilities, Eqs. (30) to (32), are built from ratios of the *γ*_*i*_, *b*_*k*_ and *y*_*l*_ to the *λ*_*i*_, so they do not depend on *A*. The immediate-return pathway, Eq. (35), visits *F, C*_1_, …, *C*_*k*_ but not *Q*, so *f* ^*R*^ is a convolution of exponentials with rates *γ*_0_, *λ*_1_, …, *λ*_*k*_, none of which involve *A*.

(ii) In Eq. (43), *A* appears only in the last term, whose coefficient is *ρ*^*C*^ + *ρ*^*S*^ = 1 − *ρ*^*R*^. The remaining terms make up *M* (*z*), which by (i) does not depend on *A*.

(iii) Differentiating 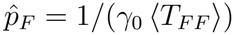 using (ii) gives Eq. (46). Its numerator clearly vanishes as *ρ*^*R*^ → 1, and its denominator is always nonzero since ⟨*T*_*F F*_ ⟩ ≥ 1*/γ*_0_ (Eq. (43)).

Let 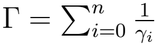. In the canonical restriction, *ρ*^*R*^ = 0 and ⟨*T*_*F F*_ ⟩ = Γ+1*/A*, so part (iii) gives

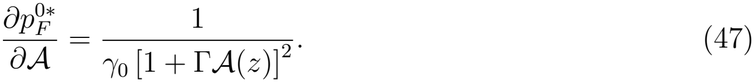

#### Corollary 1

(Selective control of the early and late portions of the return time distribution) *At fixed z, let* B_*k*_(*z*) > 0 *and ρ*^*R*^ < 1. *Then:*

*(i) β*_*k*_ *changes both the early (immediate-return) and the late (canonical and sequestration) components of the return time density, whereas A changes only the late component, ρ*^*C*^*f* ^*C*^ + *ρ*^*S*^*f* ^*S*^;

*ii) the map* 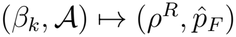 *has a triangular Jacobian with positive diagonal and is therefore locally invertible. It directly follows that tuning both parameters varies the immediate-return probability and the firing occupancy independently*.

*Proof* (i) *λ*_*k*_ depends on *b*_*k*_ = *β*_*k*_*ℬ*_*k*_, so by Eqs. (30) and (31), *ρ*^*R*^ increases and *ρ*^*C*^ decreases strictly with *β*_*k*_, while *f* ^*R*^ and *f* ^*C*^ depend on *λ*_*k*_. The statement for *A* is exactly Theorem 2(i).

ii) By (i), *∂ρ*^*R*^*/∂A* = 0 and *∂ρ*^*R*^*/∂β*_*k*_ > 0, and 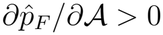 by Eq. (46).

The early hazard component *w*^*R*^*h*^*R*^ = *ρ*^*R*^*f* ^*R*^*/* (*ρ*^*R*^*S*^*R*^ + *ρ*^*C*^*S*^*C*^ + *ρ*^*S*^*S*^*S*^) depends on *A* only through *S*^*C*^ and *S*^*S*^, via neurons that have already reached *Q*, a fraction that is small when *t* is short compared with the time needed to pass from *F* to *C*_*n*_, so that *S*^*C*^(*t*) ≈ *S*^*S*^(*t*) ≈ 1. At steady state, *A* also acts through *z*^*^.

### Equivalence to Wilson–Cowan and nonlinear decay

#### Theorem 3

(Immediate return as nonlinear decay) *Imposing the steady-state relations between p*_*F*_ *and the other occupancies at all times reduces Eq. (25) to*

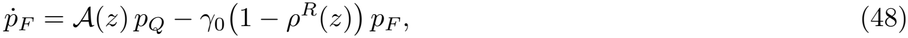

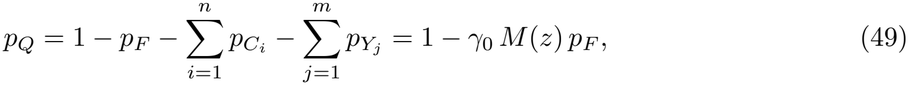

*with M as in Eq. (45), and z* = *a p*_*F*_ + *u*.

*The fixed points of this reduced system are exactly the steady states* 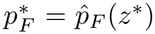, *and immediate return enters the decay of p*_*F*_ *through the effective rate γ*_eff_ (*z*) = *γ*_0_(1 − *ρ*^*R*^(*z*)), *which depends on p*_*F*_ *through z*.

*Proof* Probability flows into *F* from *Q* at rate A and from *C*_*k*_ at rate *b*_*k*_, and out of *F* to *C*_1_ at rate *γ*_0_, so the *p*_*F*_ row of Eq. (25) is exactly

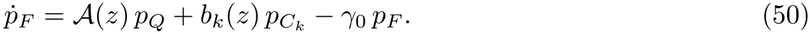

At steady state, Eqs. (27) and (8) give 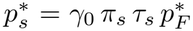 for every state. For *C*_*k*_, the probability flow through immediate return is 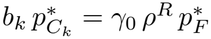. Summing over the recovery and sequestration states, with 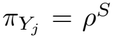 and 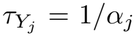, gives 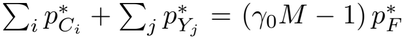. Imposing these relations at all times, with the quiescent occupancy fixed (Eq. (18)), gives Eq. (10) and Eq. (11).

Note, this reduction is only guaranteed to be exact at steady states and is only approximate elsewhere.

### Effect of sequestration

Let *T*_*sF*_ be the time from entering state *s* to the next entry into *F* . Note 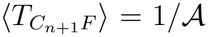, as we set *C*_*n*+1_ = *Q*. The first exit from *C*_*l*_ (the sequestration branch site) occurs after a mean sojourn time 1*/λ*_*l*_, with *λ*_*l*_ = *γ*_*l*_ + *δ*_*lk*_*b*_*k*_ + *y*_*l*_, and leads to *C*_*l*+1_, to *Y*_1_, or directly to *F* (if *l* = *k*). Conditioning on this first exit gives

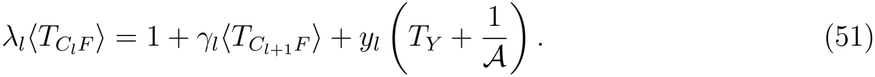

The same equation with *y*_*l*_ = 0 holds as well, where we write 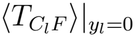. Since 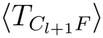 is independen)t of *y*_*l*_, subtracting the two equations gives 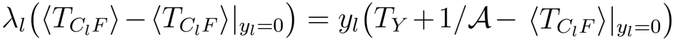 : with probability *y*_*l*_*/λ*_*l*_, sequestration replaces the remaining time to *F* by *T*_*Y*_ +1*/A*. A return reaches *C*_*l*_ with probability 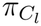, and neither 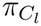 nor the time spent before reaching *C*_*l*_ depends on *y*_*l*_, so multiplying by 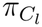 and using 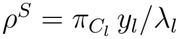 gives

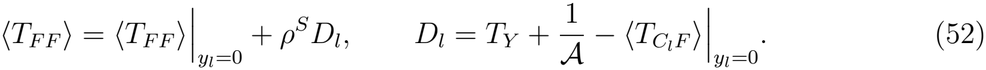

Thus, sequestration delays recovery if and only if the mean delay per event, *D*_*l*_, is positive. For sequestration at or upstream of immediate return (*l* ≤ *k*), as for the baseline parameters (*l* = 1, *k* = 2), immediate return can only shorten 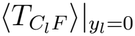 relative to its value for the canonical pathway 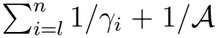, so *D*_*l*_ > 0 whenever *T*_*Y*_, the mean time to traverse the sequestration chain, exceeds 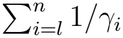, the mean time to traverse the canonical portion from *C*_*l*_ to *Q*. If immediate return lies upstream (*k* < *l*), the two values coincide and this condition is also necessary. By Eq. (27), the sequestered occupancy at steady state is 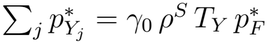, so even when *ρ*^*S*^ does not vary with input, as for *l* < *k* with *Y*_*l*_ ≡ 1, sequestration engages in proportion to firing.

### Redistribution of pathways by sequestration

#### Theorem 4

(Sequestration lowers immediate-return and raises the slower returns) *Let sequestration lie upstream of immediate-return (l* < *k) and be independent of input*, Y_*l*_ ≡ 1, *so that y*_*l*_ = *κ*_*l*_. *Let the recovery rates be constant, a* > 0 *and b*_*k*_ > 0, *and let z*^*\**^ *be a steady state at fixed u with* 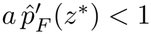 *and D*_*l*_ > 0. *Writing 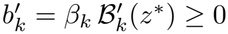 along the steady state:*

*(i) the steady-state input decreases with κ*_*l*_,

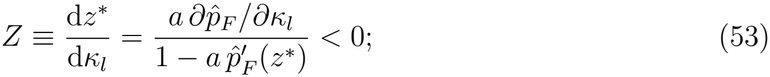

*ii) the immediate-return probability decreases and the sequestration probability increases*,

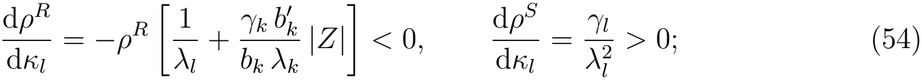

*ii) the canonical and sequestration probabilities together increase, and among returns that are not sequestered, the canonical fraction ρ*^*C*^*/*(*ρ*^*C*^ + *ρ*^*R*^) = *γ*_*k*_*/λ*_*k*_ *increases, strictly if* 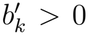. *The canonical probability itself changes as*

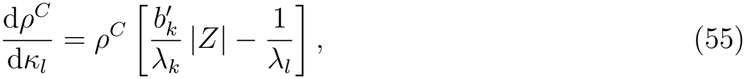

*and increases if and only if* 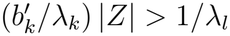.

*Proof* For *l* < *k*, Eqs. (30) to (32) read *ρ*^*C*^ = (*γ*_*l*_*/λ*_*l*_)(*γ*_*k*_*/λ*_*k*_), *ρ*^*R*^ = (*γ*_*l*_*/λ*_*l*_)(*b*_*k*_*/λ*_*k*_) and *ρ*^*S*^ = *κ*_*l*_*/λ*_*l*_, with *λ*_*l*_ = *γ*_*l*_ +*κ*_*l*_ and *λ*_*k*_ = *γ*_*k*_ +*b*_*k*_. Since Y_*l*_ ≡ 1, *κ*_*l*_ enters only *λ*_*l*_, and *z* enters only *b*_*k*_. At fixed *z*, therefore, 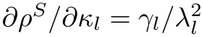, and the common factor *γ*_*l*_*/λ*_*l*_ gives *∂ρ*^*R*^*/∂κ*_*l*_ = −*ρ*^*R*^*/λ*_*l*_ and *∂ρ*^*C*^*/∂κ*_*l*_ = −*ρ*^*C*^*/λ*_*l*_. At fixed *κ*_*l*_, *ρ*^*S*^ does not depend on *z*, 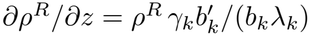 and 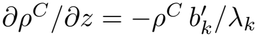.

i. *D*_*l*_ does not depend on *κ*_*l*_, so Eq. (52) gives 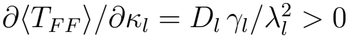 and hence 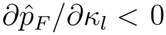. Differentiating 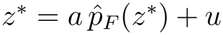 at fixed *u* gives Eq. (53), whose denominator is positive by Eq. (29), so *Z* < 0 for *a* > 0.
ii. The chain rule, d*ρ/*d*κ*_*l*_ = *∂ρ/∂κ*_*l*_ + *Z ∂ρ/∂z*, with *Z* < 0 gives Eq. (54).
iii. Since the three probabilities sum to one, d(*ρ*^*C*^ + *ρ*^*S*^)*/*d*κ*_*l*_ = −d*ρ*^*R*^*/*d*κ*_*l*_ > 0. The fraction *γ*_*k*_*/λ*_*k*_ depends on *κ*_*l*_ only through *z* and does not increase with *z*, since 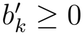, so it increases with *κ*_*l*_ when *Z* < 0, strictly if 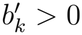. The chain rule gives Eq. (55).

Sequestration thus lowers *ρ*^*R*^ both directly and by lowering activity, raises *ρ*^*S*^ directly, and raises *ρ*^*C*^ only through activity, as in Supplementary Fig. S4. The condition for *ρ*^*C*^ to increase holds where 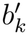 is large relative to *λ*_*k*_ and where |*Z*|, which diverges at the saddle-node, is large. It fails where *ℬ*_*k*_ saturates 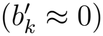 and on the unstable branch, where *Z* > 0.

### Stochastic simulations

Finite populations (Fig. 8) were simulated exactly with the Gillespie algorithm [84]. A population of *N* = 2000 neurons evolved as the joint CTMC, with neuron *i* jumping from *s* to *s*^′^ at rate 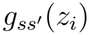 and *z*_*i*_ given by Eq. (22), and with neuron-specific parameters drawn as described in the Supplementary Material. The external drive was piecewise constant. Occupancies were compared with the homogeneous mean-field dynamics, Eq. (24). The Fano factor and the fraction of neurons firing at least once, *n*_Δ_, were computed from spike counts in windows of length Δ = 3, and each neuron’s change in firing rate was measured relative to its mean rate over *t* ∈ [10, 50).

**Table 1:**
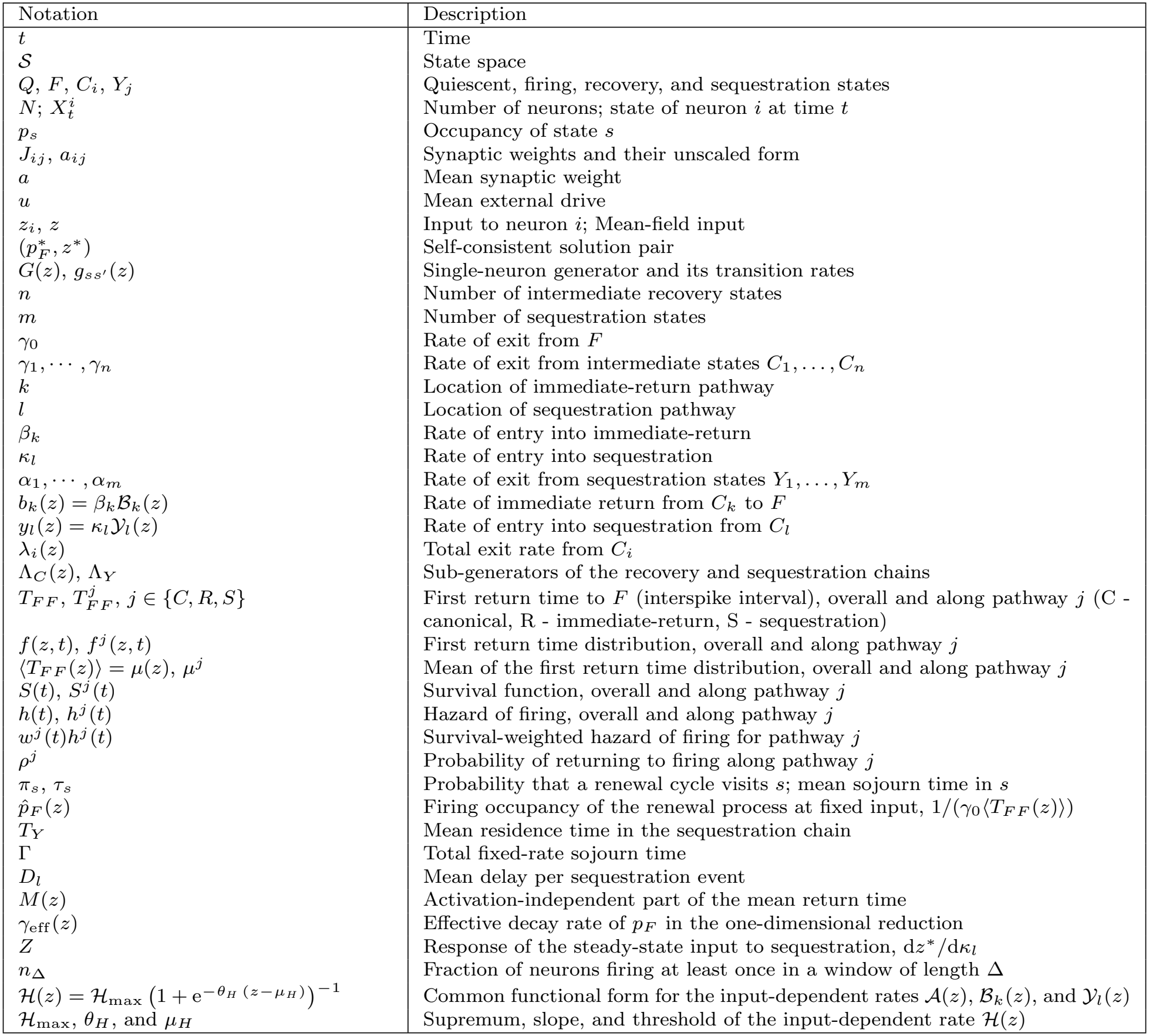
Common notations and parameters used in the model and their description.

| Notation | Description |
|---|---|
| t | Time |
| 𝒮 | State space |
| Q, F, C<sub>i</sub>, Y<sub>j</sub> | Quiescent, firing, recovery, and sequestration states |
| N; X<sub>t</sub><sup>i</sup> | Number of neurons; state of neuron i at time t |
| p<sub>s</sub> | Occupancy of state s |
| J<sub>ij</sub>, a<sub>ij</sub> | Synaptic weights and their unscaled form |
| a | Mean synaptic weight |
| u | Mean external drive |
| z<sub>i</sub>, z | Input to neuron i; Mean-field input |
| (p<sub>F</sub><sup>∗</sup>, z<sup>∗</sup>) | Self-consistent solution pair |
| G(z), g<sub>ss′</sub>(z) | Single-neuron generator and its transition rates |
| n | Number of intermediate recovery states |
| m | Number of sequestration states |
| γ<sub>0</sub> | Rate of exit from F |
| γ<sub>1</sub>, · · · , γ<sub>n</sub> | Rate of exit from intermediate states C<sub>1</sub>, . . . , C<sub>n</sub> |
| k | Location of immediate-return pathway |
| l | Location of sequestration pathway |
| β<sub>k</sub> | Rate of entry into immediate-return |
| κ<sub>l</sub> | Rate of entry into sequestration |
| α<sub>1</sub>, · · · , α<sub>m</sub> | Rate of exit from sequestration states Y<sub>1</sub>, . . . , Y<sub>m</sub> |
| b<sub>k</sub>(z) = β<sub>k</sub>ℬ<sub>k</sub>(z) | Rate of immediate return from C<sub>k</sub> to F |
| y<sub>l</sub>(z) = κ<sub>l</sub>𝒴<sub>l</sub>(z) | Rate of entry into sequestration from C<sub>l</sub> |
| λ<sub>i</sub>(z) | Total exit rate from C<sub>i</sub> |
| Λ<sub>C</sub>(z), Λ<sub>Y</sub> | Sub-generators of the recovery and sequestration chains |
| T<sub>FF</sub>, T<sub>FF</sub><sup>j</sup>, j ∈ {C, R, S} | First return time to F (interspike interval), overall and along pathway j (C - canonical, R - immediate-return, S - sequestration) |
| f(z, t), f<sup>j</sup>(z, t) | First return time distribution, overall and along pathway j |
| ⟨T<sub>FF</sub>(z)⟩ = μ(z), μ<sup>j</sup> | Mean of the first return time distribution, overall and along pathway j |
| S(t), S<sup>j</sup>(t) | Survival function, overall and along pathway j |
| h(t), h<sup>j</sup>(t) | Hazard of firing, overall and along pathway j |
| w<sup>j</sup>(t)h<sup>j</sup>(t) | Survival-weighted hazard of firing for pathway j |
| ρ<sup>j</sup> | Probability of returning to firing along pathway j |
| π<sub>s</sub>, τ<sub>s</sub> | Probability that a renewal cycle visits s; mean sojourn time in s |
| p̂<sub>F</sub>(z) | Firing occupancy of the renewal process at fixed input, 1/(γ<sub>0</sub>⟨T<sub>FF</sub>(z)⟩) |
| T<sub>Y</sub> | Mean residence time in the sequestration chain |
| Γ | Total fixed-rate sojourn time |
| D<sub>l</sub> | Mean delay per sequestration event |
| M(z) | Activation-independent part of the mean return time |
| γ<sub>eff</sub>(z) | Effective decay rate of p<sub>F</sub> in the one-dimensional reduction |
| Z | Response of the steady-state input to sequestration, dz<sup>∗</sup>/dκ<sub>l</sub> |
| n<sub>Δ</sub> | Fraction of neurons firing at least once in a window of length Δ |
| $\mathcal{H}(z) = \mathcal{H}_{\max}\left(1 + \mathrm{e}^{-\theta_H\,(z-\mu_H)}\right)^{-1}$ | Common functional form for the input-dependent rates 𝒜(z), ℬ<sub>k</sub>(z), and 𝒴<sub>l</sub>(z) |
| ℋ<sub>max</sub>, θ<sub>H</sub>, and μ<sub>H</sub> | Supremum, slope, and threshold of the input-dependent rate ℋ(z) |

### Numerical methods and parameters

All input-dependent rates (except *Y*_*l*_ ≡ 1 for pharmacological sequestration) share the sigmoidal form used previously [62],

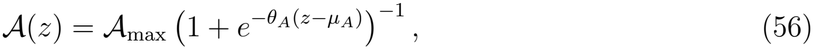

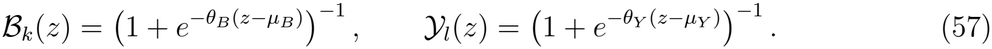

In Fig. 3(f), *A*(*z*) is replaced by *A*(*z*^*ν*^). Steady states and their stability were obtained from Eqs. (28) and (29). All analyses and simulations were implemented in MATLAB R2024b.

## Data availability

The in vivo whole-cell recordings re-analyzed in this study were reported by Otomo et al. [32] and are publicly available from the Harvard Dataverse (doi:10.7910/DVN/Q9ZXLZ). All other data were generated by numerical simulation of the model described in the Methods.

## Code availability

The code used to analyze the recordings and to simulate the model is available at https://github.com/Aero264/hazardhyperexcitability.

## Acknowledgments

We thank Profs Derk-Jan Dijk and Anne Skeldon for their support. AKK is supported by the UK Dementia Research Institute [UK DRI-7206 and PB3197] through UK DRI Ltd, principally funded by the UK Medical Research Council.

## Author Contributions

Conceptualization: AKK; Formal analysis: AKK & MPS; Validation: AKK & MPS; Writing — Original draft: AKK & MPS; Writing — Review and Editing: AKK & MPS; Funding acquisition: AKK.

## Declaration of generative AI and AI-assisted technologies in the manuscript preparation process

During the preparation of this work, the authors used ChatGPT (OpenAI) and Claude (Anthropic) to improve clarity and readability of the manuscript text, and to check the consistency of mathematical notation. The authors also used these tools to review the analysis code for potential errors, minimize its dependencies, and streamline its documentation. All scientific content, analyses, interpretations, and conclusions were developed by the authors. The authors reviewed and edited all AI-assisted material and take full responsibility for the content of the published article.

