## Supplementary Material for "Competing routes back to firing shape population hyperexcitability and its selective suppression"

### Analysis of excluded neurons from dopamine neuron recordings

In the main text, we re-analyzed *in vivo* whole-cell recordings of spontaneously active midbrain dopamine neurons from Otomo et al. [1]. For each neuron, spikes were detected using the method published in Otomo et al. [1]. Interspike intervals (ISIs) were calculated as the differences in time between successive spike peaks. $V_{\mathrm{min}}$ was the minimum of the filtered trace between two successive spike peaks. We then tested whether the ISI hazard is monotone, divided the intervals into distinct classes by the trough voltage $V_{\mathrm{min}}$ preceding each spike, and decomposed the pooled hazard into class-specific contributions.

Neurons were assigned to the monotone group if the test for a monotone hazard did not reject monotonicity and $V_{\mathrm{min}}$ had a single class (59 cells), and to the non-monotone group if the test rejected monotonicity, $V_{\mathrm{min}}$ had two classes, and these classes passed the permutation test of the class decomposition (22 cells). All other cells were excluded from group comparisons presented in the main text. Here, we present examples of neurons that were excluded.

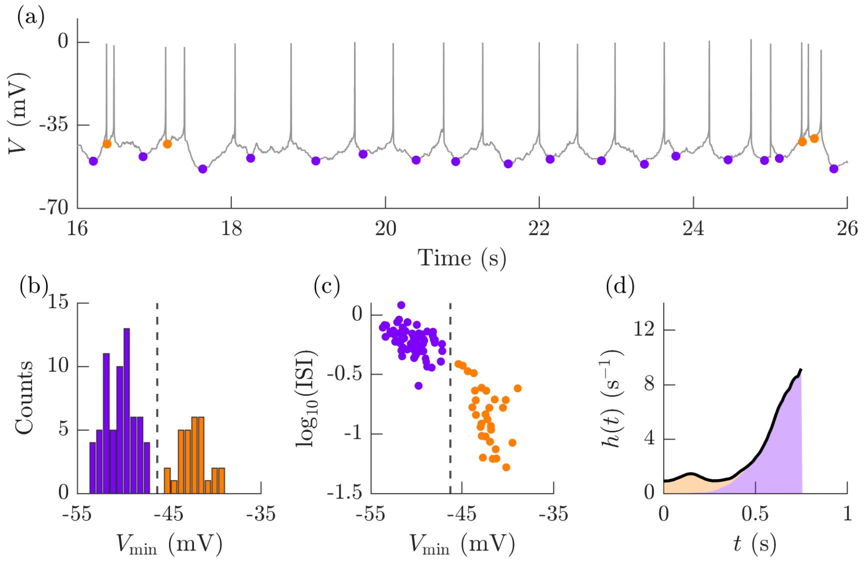


**Fig. S1**: **Example neuron with two-class $V_{\mathrm{min}}$ but passed the test for a monotone hazard.** (a) Time course of membrane potential ($V$), with $V_{\mathrm{min}}$ preceding each spike marked. Both solitary spikes and burst-initiating spikes follow deep hyperpolarized troughs (purple), whereas subsequent intraburst spikes follow more depolarized troughs (orange). (b) The distribution of $V_{\mathrm{min}}$ resolves into two distinct classes. (c) The joint distribution of $V_{\mathrm{min}}$ and $\log_{10}$(ISI). (d) The corresponding aggregate hazard shows a small dip and is resolvable into two distinct components. But the test for a monotone hazard did not reject monotonicity as the dip is much smaller and is of the order of the excursions from monotonicity that arise due to noise.

Figure S1 shows an example of a neuron that admitted a two-class decomposition of $V_{\text{min}}$ but was excluded because it passed the test for a monotone hazard because the aggregate hazard did not exhibit excursions from monotonicity of sufficient magnitude. A total of four neurons belonged to this exclusion class.

Figure S2 shows an example of the opposite case. Here, the neuron failed the monotonicity test but $V_{\text{min}}$ cold be resolved into two classes as the hyperpolarized troughs preceding the solitary spikes and the burst initiating spikes are not easily distinguishable from the relatively depolarized troughs preceding subsequent spikes within a burst. A total of eleven neurons belonged to this exclusion class.

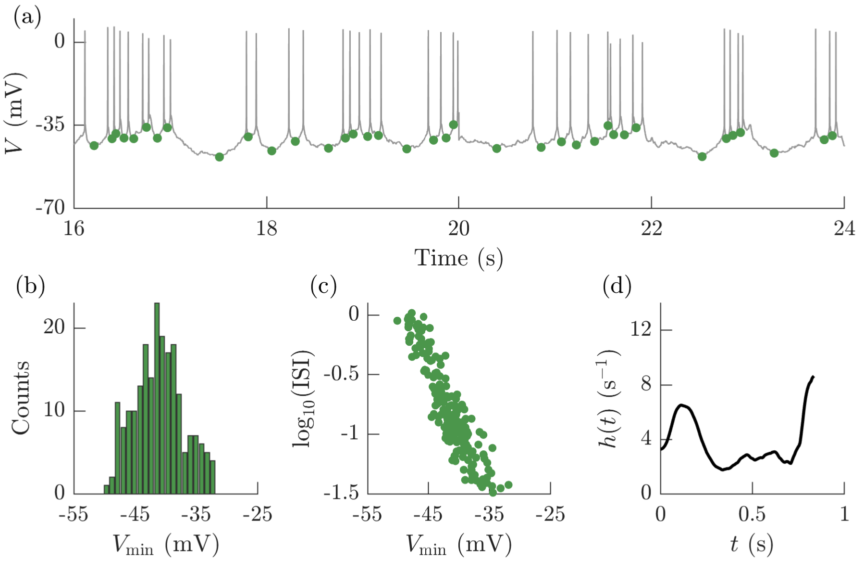


**Fig. S2**: **Example neuron that failed the test for a monotone hazard but has one-class $V_{\text{min}}$.** (a) Time course of membrane potential ($V$), with $V_{\text{min}}$ preceding each spike marked in green. The trace shows predominantly solitary spikes with intermittent bursting. (b) The distribution of $V_{\text{min}}$ forms a single cluster as the hyperpolarized troughs preceding the solitary spikes and the burst initiating spikes are not easily distinguishable from the relatively depolarized troughs preceding subsequent spikes within a burst. (c) The joint distribution of $V_{\text{min}}$ and $\log_{10}$(ISI) also forms a single cluster. (c) The corresponding hazard shows non-monotonicity rising sharply first, then falling and then rising again towards the tail. Even if the tail peak were to be attributed to end effects, the preceding fall is sufficient to rule out a monotonically increasing aggregate hazard.

### Immediate-return pathway provides a control dimension on population activity distinct from activation

In the main text, we demonstrated that the parameters that influence rates downstream of $C_k$ in the canonical pathway — including those of the activation function — do not affect neurons that return to $F$ directly from $C_k$ along the immediate-return pathway (Theorem 2). We illustrated this with the example of the rate of immediate-return $\beta_k$ and the supremum of the activation function $\mathcal{A}_{\text{max}}$ in Figure 5 of the main text. Here, we show that this holds for other parameters such as threshold and slope (Fig. S3).

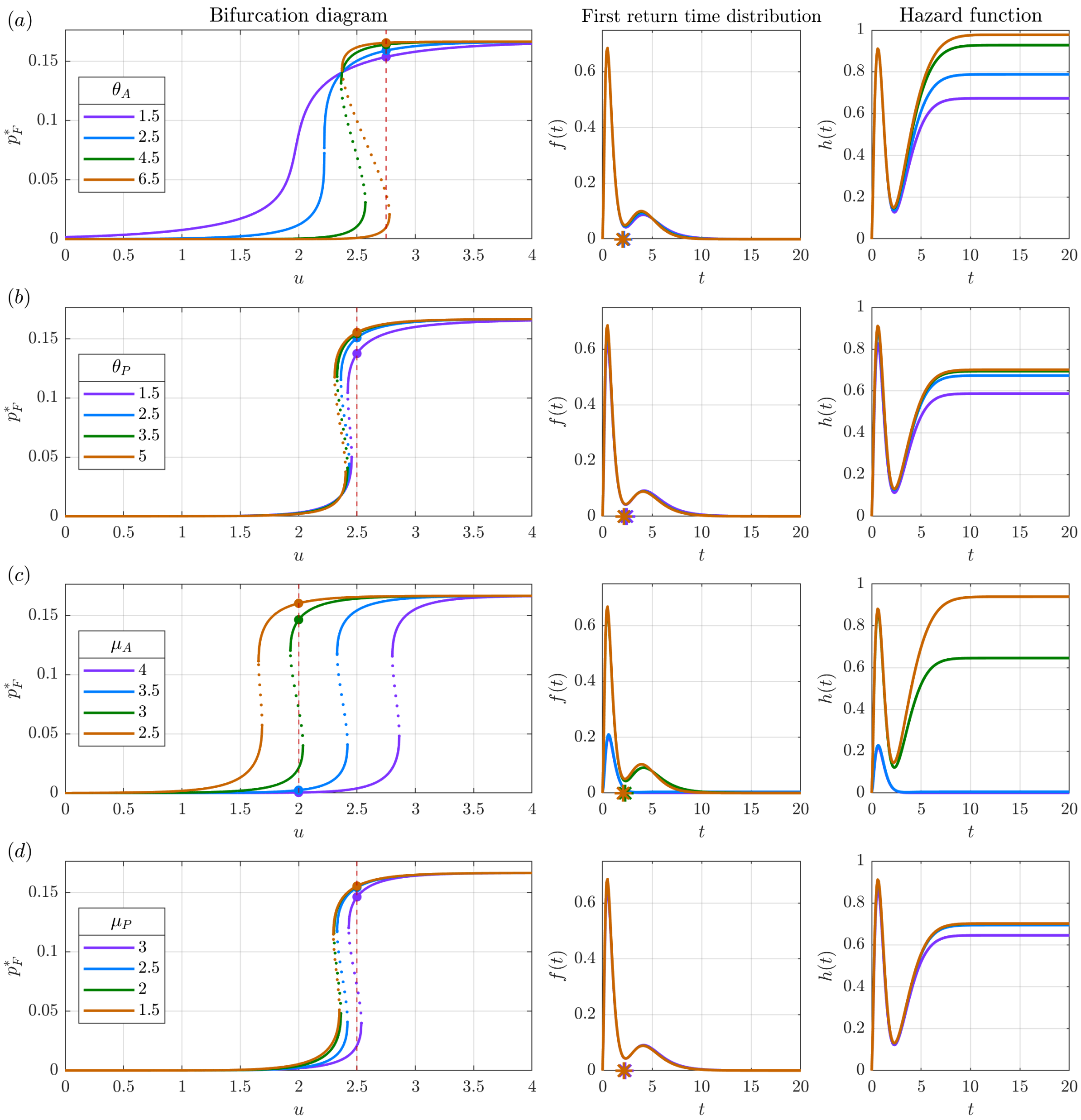


**Fig. S3**: **Perturbing canonical parameters increases the late hazard while perturbing immediate-return parameters increases the early hazard, providing independent tuning.** Variation in four different parameters lead from a monostable low-activity attractor to a monostable high-activity attractor, with an intervening bistable regime for a subset of parameter values: (a) Increasing the slope of the activation function $\theta_A$, or (c) lowering its threshold $\mu_A$ increase firing occupancy by increasing the late hazard while leaving the early hazard unchanged. Increasing the equivalent quantities for immediate-return, (b) the slope $\theta_P$, and (d) the threshold $\mu_P$ increase firing occupancy by increasing both the early and late hazard. In each case, the left panels show the 1-D bifurcation diagram for the variation in the firing occupancy $p_F^*$ against external drive $u$. Stable branches are indicated in solid lines while the unstable branch is indicated in dotted line. For select points indicated in the bifurcation diagram, the middle and the right panels show the first return time distribution $f(t)$, with its mean marked as an asterisk, and the hazard function $h(t)$.

### Sequestration enables selective suppression of pathological activity

Here, we illustrate Theorem 4 of the main text, where increasing the rate of sequestration $\kappa_l$ lowers the proportion of neurons returning to firing through the immediate-return pathway and raises the slower returns through the canonical and sequestration pathways (Fig. S4).

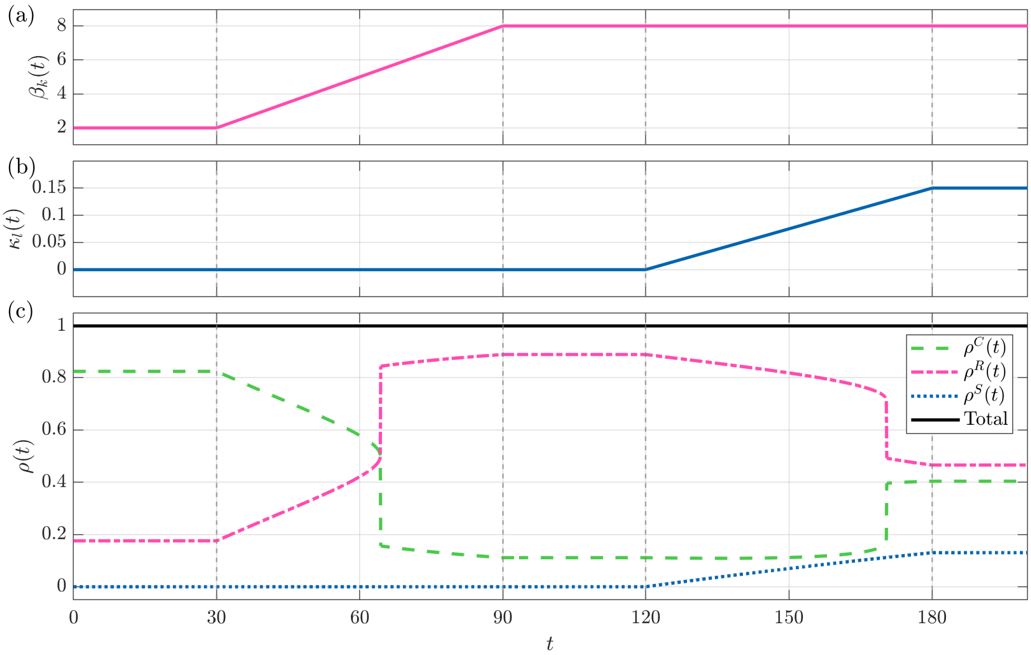


**Fig. S4: Increasing the rate of sequestration selectively diverts neurons away from the immediate-return pathway, leading to the suppression of pathological hyperexcitability.** In the figure, the trajectory follows the progression represented by panels (b-d), then (h-e) of Fig. 6 of the Main text: from untreated physiological bursting to pathological high activity, followed by drug-treated low activity resembling physiological bursting. The population is initiated at a low-activity attractor corresponding to physiological bursting with $\beta_k = 2$ and $\kappa_l = 0$. Holding all other parameters constant, $\beta_k$ is quasi-statically increased from two to eight, with the low-activity attractor losing stability and the population settling onto the high-activity attractor at $\beta_k \approx 3.8$. This is marked by a sharp increase in the proportion of neurons taking the immediate-return pathway. Subsequently, increasing $\kappa_l$ diverts some of those neurons taking the immediate-return onto the sequestration pathway, while the proportion of neurons taking the canonical pathway remains approximately constant. This diversion destabilizes the high-activity attractor and the population returns to the low-activity attractor at $\kappa_l \approx 0.126$. The annihilation of the high-activity attractor is marked by a sharp decrease in $\rho^R$.

### Parameters used for simulation

**Table 1: Parameters used for the simulation of Figs. 3 to 7 of the Main Text and Figs. S3 and S4 of the Supplementary Material.**

| Parameter | Fig. 3 | Figs. 4, 5 and S3 | Figs.6, 7 and S4 |
|---|---|---|---|
| $a$ | 8 | 8 | 8 |
| $u$ | – | 2.4 | 1.75 |
| $n$ | 10 | 10 | 10 |
| $\gamma_0$ | 3 | 3 | 1 |
| $\gamma_1 = \cdots = \gamma_n = \gamma$ | 3 | 3 | 1 |
| $k$ | – | 2 | 2 |
| $\beta_k$ | – | 6 | – |
| $l$ | – | – | 1 |
| $\alpha_1 = \cdots = \alpha_m = \alpha$ | – | – | 1 |
| $m$ | – | – | 30 |
| $\mathcal{A}_{\max}$ | 1 | 1 | 1 |
| $\mu_A$ | 3.5 | 3.5 | 3.5 |
| $\theta_A$ | 3.5 | 3.5 | 2.5 |
| $\mathcal{B}_{\max}$ | – | 1 | 1 |
| $\mu_B$ | – | 2.5 | 2.5 |
| $\theta_B$ | – | 3.5 | 3.5 |

**Table 2: Quenched heterogeneity parameters used for the simulation of Fig. 8 of the Main Text.** For log-normal distribution, the column "Variation" indicates the coefficient of variation whereas for Gaussian distribution, it indicates the standard deviation.

| Parameter | Mean | Variation | Distribution |
|---|---|---|---|
| $\gamma_0$ | 3 | 0.2 | log-normal |
| $\gamma_1 = \cdots = \gamma_n = \gamma$ | 3 | 0.2 | log-normal |
| $\alpha_1 = \cdots = \alpha_m = \alpha$ | 3 | 0.2 | log-normal |
| $\beta_k$ | 6 | 0.3 | log-normal |
| $\kappa_l$ | 1 | 0.3 | log-normal |
| $\theta_A$ | 2 | 0.1 | log-normal |
| $\theta_B$ | 3.5 | 0.1 | log-normal |
| $\theta_Y$ | 5.5 | 0.1 | log-normal |
| $\mu_A$ | 3 | 0.05 | Gaussian |
| $\mu_B$ | 2.5 | 0.05 | Gaussian |
| $\mu_Y$ | 8 | 0.15 | Gaussian |

**Table 3: Other fixed parameters used for the simulation of Fig. 8 of the Main Text.**

| Parameter | $a$ | $n$ | $m$ | $k$ | $l$ | $\mathcal{A}_{\max}$ | $\mathcal{B}_{\max}$ | $\mathcal{Y}_{\max}$ |
|---|---|---|---|---|---|---|---|---|
| Value | 8 | 10 | 50 | 2 | 1 | 1 | 1 | 1 |
